# Transcriptomic landscape of mammalian ventral pallidum at single-cell resolution

**DOI:** 10.1101/2024.05.24.595793

**Authors:** Lite Yang, Lisa Z Fang, Michelle R Lynch, Chang S Xu, Hannah Hahm, Yufen Zhang, Monique R Heitmeier, Vincent Costa, Vijay K Samineni, Meaghan C Creed

**Author notes:** Corresponding Authors: Meaghan C Creed –, Vijay K Samineni.

## Abstract

The ventral pallidum (VP) is critical for motivated behaviors. While contemporary work has begun to elucidate the functional diversity of VP neurons, the molecular heterogeneity underlying this functional diversity remains incompletely understood. We used snRNA-seq and *in situ* hybridization to define the transcriptional taxonomy of VP cell types in mice, macaques, and baboons. We found transcriptional conservation between all three species, within the broader neurochemical cell types. Unique dopaminoceptive and cholinergic subclusters were identified and conserved across both primate species but had no homolog in mice. This harmonized consensus VP cellular atlas will pave the way for understanding the structure and function of the VP and identified key neuropeptides, neurotransmitters, and neuro receptors that could be targeted within specific VP cell types for functional investigations.

**Teaser:** Genetic identity of ventral pallidum cell types is conserved across rodents and primates at the transcriptional level

## Introduction

The ventral pallidum (VP) is a hub within the ventral basal ganglia. The VP is the primary output of the ventral striatum and orchestrates activity of midbrain dopamine neurons and thalamic nuclei through direct and polysynaptic connections (*1*). The VP has been proposed to play the role of a ‘limbic motor interface’, regulating conditioned responses to appetitive and aversive stimuli. In rodents, manipulations of VP cellular activity produce profound effects on drug self-administration (*2–4*), decision making under conflict (*5–9*) and feeding (*10*, *11*). In humans, the VP is activated in response to reward-predictive cues (*12*) and the magnitude of this activation tracks the subjective and state-dependent value of rewards (*13*). Together, this points to the VP as a tractable brain target for neuromodulation therapies aimed at treating disorders of dysfunctional reward processing.

The VP was considered to be comprised primarily of inhibitory GABAergic neurons and neurons belonging to the basal forebrain cholinergic system (*14*). Contemporary circuit dissection, however, has revealed the VP also contains excitatory glutamatergic neurons. The projections of these glutamatergic neurons follow the same pathways as prototypical GABAergic neurons, but constitute a discrete population (*6*, *7*). These two populations, coarsely defined by whether they release glutamate or GABA, exert opposing effects on appetitive behavior with glutamatergic neurons constraining and GABAergic neurons promoting pursuit of rewards. Despite this functional heterogeneity, the molecular diversity within each of these neurochemically-defined subclasses remains poorly understood. Finally, our current understanding of the VP cellular makeup has been dissected along the expression of genes encoding transcription factors, calcium-binding proteins, or neurotransmitter receptors (i.e. parvalbumin [*Pvalb*], dopamine receptor D3 [*Drd3*] or neuronal PAS domain protein 1 [*Npas1*], *9*, *15*, *16*). Each of these broadly-defined populations was anatomically, biophysically, and functionally heterogeneous. Though these studies revealed valuable functional insights into the VP, they do not fully capture the cellular taxonomy of VP neurons. Thus, an overarching framework for understanding the molecularly-defined cell types within the VP, and thus for integrating these prior circuit-level studies into an overarching model of VP function, is lacking.

Given its role in modulating reward evaluation, seeking, consumption, and associative learning, many circuit dissection studies have proposed the VP to be a target for therapies for mental and emotional disorders. A key first step towards translation is determining whether cell types under investigation in rodents are conserved in primate species. As in rodent studies (*5*, *17–21*), *in vivo* recording in primates during reward-guided choice assays reveals that response profiles of VP neurons to rewards and reward-predictive cues are heterogeneous, and VP neuronal firing reflects discrete attributes of rewards and operant responding (*22–25*). However, the cellular diversity of the VP has not been characterized in recent transcriptomic atlases of the macaque (*26*) and marmoset brain (*27*). Given the difficulty in obtaining high-quality human samples and the evolutionary proximity of non-human primates (NHPs) to humans, old world monkeys are an ideal model organism for understanding principles governing transcriptomic profiles of basal ganglia neurons. Here, we performed single-nucleus RNA sequencing (snRNA-seq) of the VP across one rodent and two primate species, macaques (*Macaca mulatta*) and olive baboons (*Papio anubis*), followed by spatial validation of identified VP cell types. The transcriptional taxonomy presented here emphasizes the conserved nature of VP cell types across mammalian species and lays a foundation for understanding how the activity of these cell types modulates reward-guided behavior.

## Results

### Mouse atlas of ventral pallidum

To profile the transcriptional diversity of VP neurons, we performed snRNA-seq on four VP samples (two female and two male samples, Fig. 1A, Methods). After quality control and removal of low-quality nuclei and doublets, 38,130 nuclei were retained in the final dataset with an average of 2,105.7 genes per nucleus (Fig. S1A, Table S1). Notably, nuclei from individual libraries were represented in each cluster, indicating a low level of technical variation (Fig. S1B). Clustering analysis revealed eight major transcriptional cell types including neurons (65.2% of the total nuclei), astrocytes (10.8%), choroid epithelial cells (0.2%), macrophages (0.1%), microglia (4.5%), oligodendrocytes (12.3%), oligodendrocyte progenitor cells (OPC, 5.4%), and vascular cells (1.5%; Fig. 1B). These cell types exhibit distinct transcriptional profiles and are marked by high expression levels of canonical marker genes of each type (Fig. 1C, Table S2;(*28*, *29*).

**Fig. 1.**
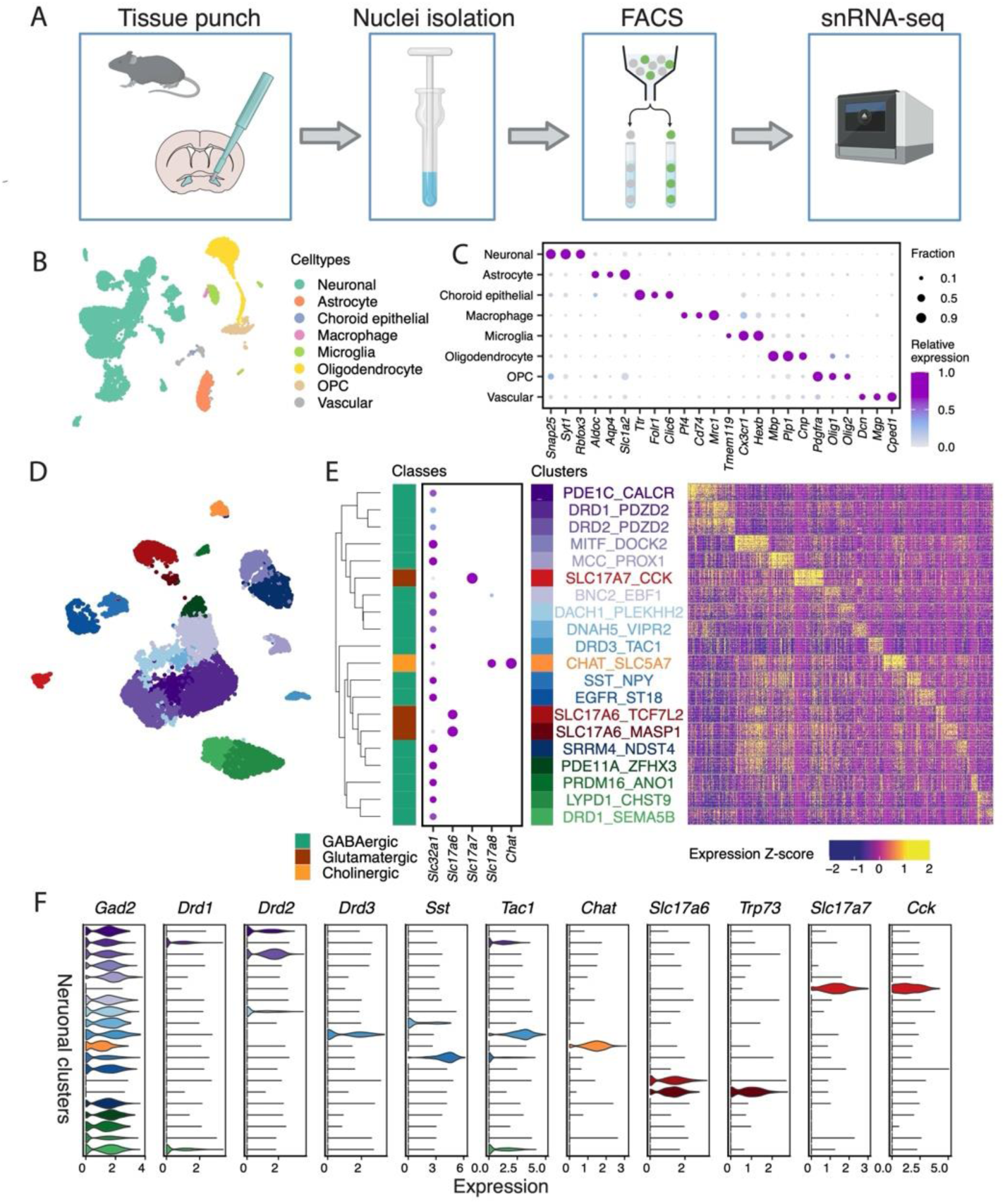
An atlas of the mouse ventral pallidum is revealed by snRNA-seq. (**A**) Experimental workflow of snRNA-seq of mouse VP. (**B**) Uniform manifold approximation and projection (UMAP) visualization of 10,000 downsampled nuclei, colored by cell types. (**C**) Dot plot displaying the expression of select cell-type-specific marker genes in individual cell types. Dot size denotes the fraction of nuclei expressing a marker gene (>0 counts), and color denotes relative expression of a gene in each cell type (calculated as the mean expression of a gene relative to the highest mean expression of that gene across all cell types). (**D**) UMAP of 10,000 downsampled mouse neuronal nuclei, colored by neuronal clusters. (**E**) Left: hierarchical clustering of neuronal clusters based on top 100 marker genes (sorted by Log2FC) per cluster. Middle: Dot plot displaying the expression of select neurotransmitter genes for class annotation. See Fig 1C for legend. Right: heatmap showing Z-scores of the expression of top 100 marker genes (sorted by Log2FC) per neuronal clusters in individual nuclei of snRNAs-seq. 50 nuclei are downsampled per neuronal cluster and displayed on the heatmap. (**F**) Violin plot showing log-transformed expression of select marker genes in individual neuronal clusters. Neuronal clusters are ordered according to the dendrogram in E.

Among all transcriptional cell types, neurons exhibited a high degree of transcriptional heterogeneity based on inspection of the UMAP (Fig. 1B). Unsupervised clustering of the neuronal nuclei revealed 20 neuronal clusters. Sixteen of these clusters are GABAergic (90.1% of neuronal nuclei) with high expression of *Slc32a1* encoding vesicular GABA transporter (VGAT). They also exhibit distinct canonical marker genes such as *Gad1*, and *Gad2*. There are four clusters of glutamatergic neurons (9.9%) labeled with high expression of *Slc17a6*, *Slc17a7, or Slc17a8* encoding vesicular glutamate transporter (VGLUT) isoforms 2, 1, and 3, respectively (Fig. 1D,E, Fig S1C). One glutamatergic cluster is also cholinergic (1.5%) based on the expression of *Chat*.

While most nuclei exhibit distinct expression of only one of the neurochemical class marker genes, 0.95% of neuronal nuclei co-express *Slc32a1* with one of the vesicular glutamate transporters, and four clusters, including the cholinergic cluster, are highly enriched with double-positive nuclei (2.4 – 7.2% of total nuclei in each of the four clusters, Fig. S1D). This observation is unlikely due to technical artifacts as the double positive populations are still present after ambient RNA correction (Fig. S1E). Within each of those classes, neuronal clusters exhibit distinct transcriptional profiles and thus we named them by a combination of marker genes that uniquely labeled each cluster (Fig. 1E, Fig. S1C, Table S2, Methods).

Among the 16 clusters of GABAergic neurons identified in the mouse VP, many corresponded to the GABAergic neurons previously described in basal forebrain structures (*28*, *30*, *31*). GABAergic neuronal clusters were differentiated based on the enrichment of genes encoding neuropeptides (i.e. *Sst*, *Tac1, Vip*), neurotransmitter receptors (i.e. *Calcr*, dopamine receptors, *Drd1*, *Drd2*, *Drd3)*, and/or transcription factors implicated in cell fate determination and neural differentiation (i.e. *Lypd1*, *Dach1*, *Mitf*, *Prox1*, *Ebf1*, *Zfhx3*, *Prdm16*, *Srrm3*). Three clusters of GABAergic neurons (PDE1C_CALCR, DRD1_PDZD2, AND DRD2_PDZD2) were hierarchically organized into dendrograms based on transcriptional similarity and characterized by the highest expression level among all neuronal clusters of *Pdzd2*, a gene encoding for a PDZ-domain containing protein previously used as a marker of striatal cell types (*28*). Using fluorescent in situ hybridization (FISH), we confirmed the expression of *Pdzd2* throughout VP territories ventral to the anterior commissure, arguing against the classification of *Pdzd2*-expressing neurons as a selective striatal marker (Fig. S2; *28*, *29*).

We also characterized four glutamatergic clusters (based on the expression of one of the three isoforms of VGLUT), one of which represents neurons of the basal forebrain cholinergic system (CHAT_SLC5A7), with distinct expression *Slc17a8* (VGLUT3) as previously reported (Fig. S1C)(*34*). We also identified a sparse but discrete population of *Slc17a7* (VGLUT1)-expressing neurons (SLC17A7_CCK), which we confirmed with FISH (Fig. S2). *Slc17a7* was highly co-localized with *Cck,* which encodes neuropeptide cholecystokinin, analogous to neuronal subpopulations previously described in the extended amygdala (*35*, *36*). Finally, our clustering revealed two distinct populations of *Slc17a6* (VGLUT2)-expressing neurons in the mouse VP (SLC17A6_TCF7L2 and SLC17A6_MASP1; Fig. 1D,E). Substantial prior work has demonstrated these *Slc17a6*-positive VP neurons are largely non-overlapping with GABAergic populations (*6*, *7*), which we confirm. Our results comport with publicly available MERFISH datasets (*30*, *31*) confirming the presence of two *Slc17a6*-positive neuronal populations throughout the basal forebrain. *Slc17a6*-positive VP neurons were differentiated by their expression of *Trp73*, which was expressed exclusively in the minor *Slc17a6*-positive cluster SLC17A6_MASP1 (Fig. 1F) (*28*, *29*).

VP activity is necessary for sex-specific parenting behavior (*37*), and sex differences in peptidergic inputs to the VP have been reported (*38*). Therefore, we next asked if the VP showed sex-specific differences in cell type composition and gene expression. Analysis revealed that male and female VPs contain broadly the same cell types (Fig. S1F) that express highly similar marker genes (Pearson’s r = 0.92 - 0.99 per neuronal cluster/cell type between males and females (Fig. S1G). To further investigate sex-specific gene expression in VP cell types, we next identified genes differentially expressed in nuclei from male and female samples in distinct VP neuronal clusters or non-neuronal cell types (Methods). The most dramatic sex differences across cell types are known sex-specific genes that are more highly expressed in females and involved in X chromosome inactivation (e.g., *Xist*, *Tsix*) or Y chromosome genes (e.g., *Uty*, *Ddx3y, Eif2s3y, Kdm5d*) that are more highly expressed in males (Fig. S1H). In addition to known sex-specific gene expression, we also identified six genes with sex-specific expression in individual cell types (Fig S1I), which may potentially underlie sex-specific VP functions.

### Spatial validation of select transcriptional clusters

To corroborate the expression of marker genes for select VP neuronal clusters and to visualize the spatial location of these clusters across the VP, we performed HiPlex FISH in mouse tissue. We prioritized canonical VP marker genes that also distinctly label GABAergic (Fig 2) and non-GABAergic (Fig 3) subsets of VP neurons. In total, 24,477 nuclei were segmented from images taken across four coronal sections representing the anterior to posterior extent of the VP, with 4,631 of those nuclei being positively labelled by at least one probe. The anatomical coordinates of nuclei were then reconstructed and compared to our snRNA-seq results to corroborate the expression of the identified cell types in the VP. Consistent with the snRNA-seq, *Gad2*-positive nuclei are highly enriched in the VP, *Slc17a6*-positive nuclei are sparser, with only 1.22% of all *Slc17a6*-positive cells co-expressing *Gad2*, consistent with prior work (*6*, *7*).

**Fig. 2.**
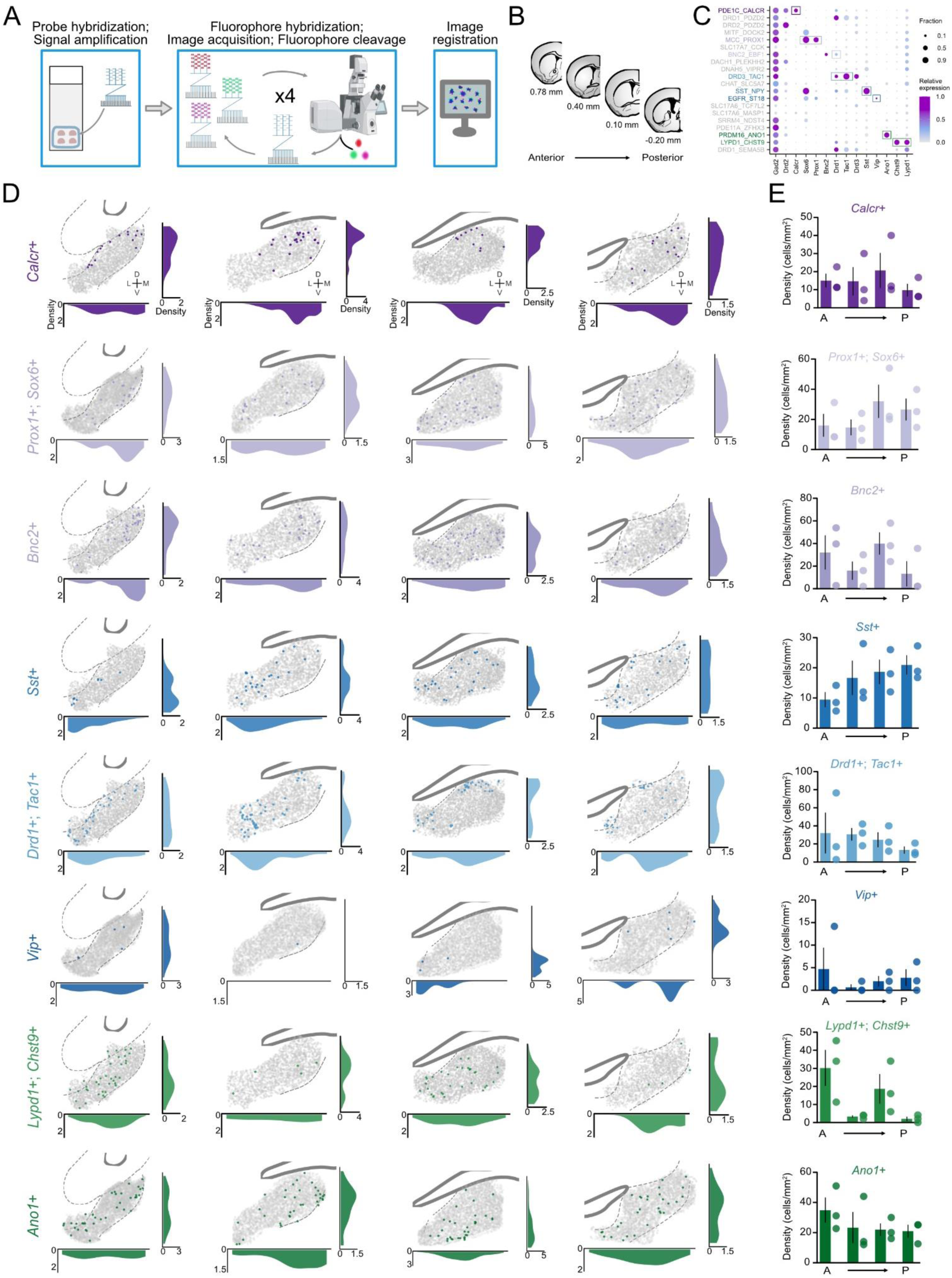
Multiplexed in situ hybridization corroborates spatial organization of select GABAergic neuronal clusters in mouse VP. (**A**) Experimental workflow of HiPlex FISH in mouse VP. (**B**) Diagram depicting the approximate anteroposterior levels of the VP that were assessed. (**C**) Dot plot displaying the expression of gene markers that were used to selectively mark each cluster. Unique gene marker(s) and corresponding cluster name in parentheses are as follows: *Calcr* (PDE1C_CALCR); co-expression of *Prox1* and *Sox6* (MCC_PROX1); *Bnc2* (BNC2_EBF1); co-expression of *Drd1* and *Tac1* (DRD3_TAC1); *Sst* (SST_NPY); EGFR_ST18 (*Vip*); co-expression of *Lypd1* and *Chst9* (LYPD1_CHST9), and *Ano1* (PRDM16_ANO1). (**D**) Spatial maps of cells expressing *Calcr*, *Prox1* and *Sox6*, *Bnc2*, *Drd1* and *Tac1*, *Sst*, *Vip*, *Lypd1* and *Chst9*, and *Ano1* over four levels of the VP. Each map is integrated over all biological samples (n=3), with colorized dots representing cells that expressed specified marker genes, and gray dots representing cells that did not express these marker genes. Density plots depict the relative distribution of positively-identified cells in respective clusters over the mediolateral and dorsoventral extent of the VP. (**E**) Bar graphs indicating the density of positively-identified cells belonging to individual clusters per mm^2^ over the anteroposterior axis of the VP.

**Fig. 3.**
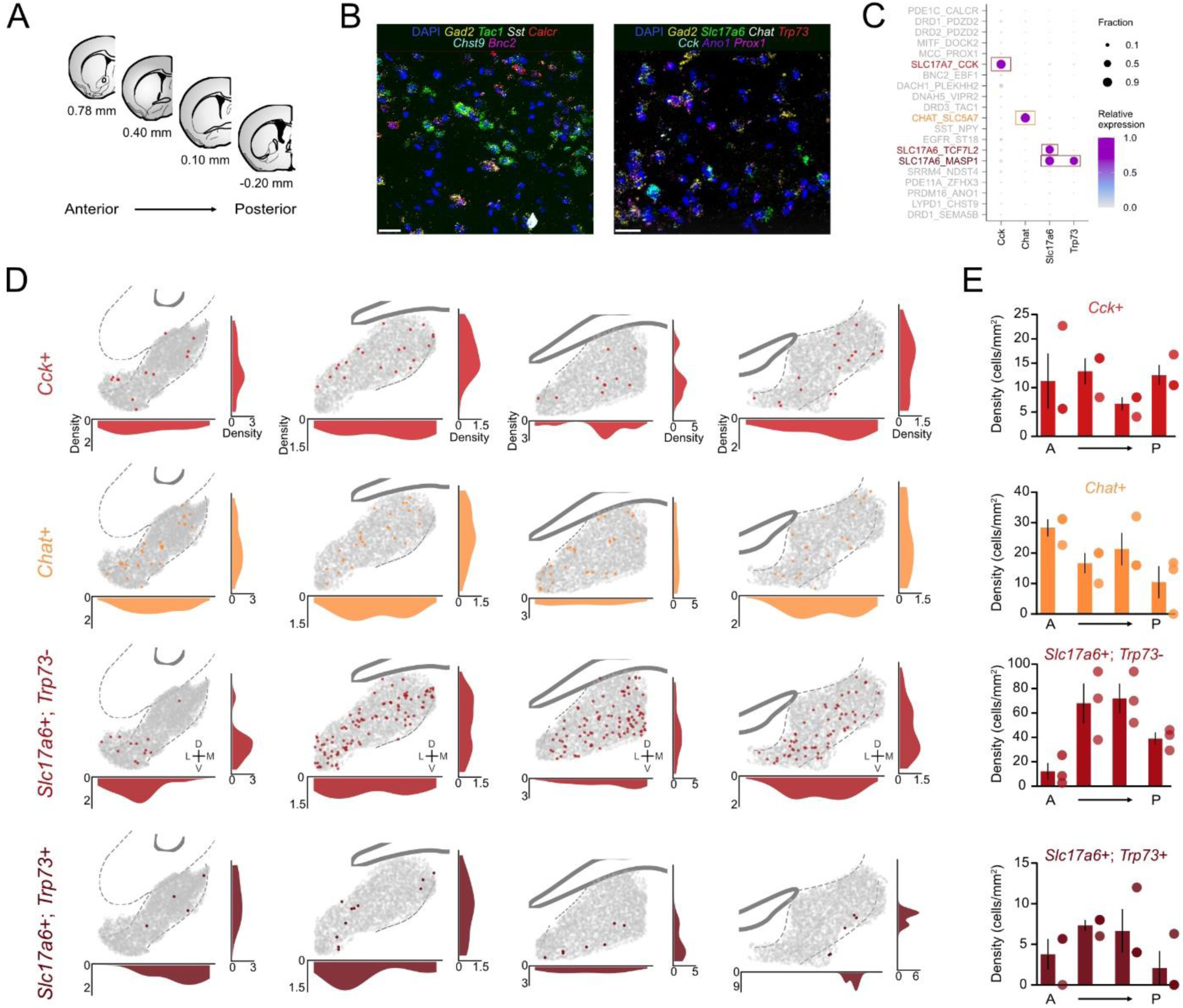
Multiplexed in situ hybridization corroborates spatial organization of select non-GABAergic neuronal clusters in mouse VP. (**A**) Diagram depicting the approximate anteroposterior levels of the VP that were assessed. (**B**) Representative HiPlex FISH images showing select probes for marker genes used to differentiate specific neuronal clusters. DAPI was used as a counterstain. Probes were hybridized *to Gad2, Tac1, Sst, Calcr, Chst9, Bnc2*, *Slc17a6*, *Chat*, *Trp73*, *Cck*, *Ano1*, and *Prox1* mRNA. Scale bar = 20 µm. (**C**) Dot plot displaying the expression of gene markers that were used to selectively mark each cluster. Unique gene marker(s) and corresponding cluster name in parentheses as follows: *Cck* (SLC17A7_CCK); *Chat* (CHAT_SLC5A7); *Slc17a6* without *Trp73* (SLC17A6_TCF7L2); co-expression of *Slc17a6* and *Trp73* (SLC17A6_MASP1). (**D**) Spatial maps of cells expressing *Cck*, *Chat*, *Slc17a6* without *Trp73*, and *Slc17a6* with co-expression of *Trp73* over four levels of the VP. Each map is integrated over all biological samples (n=3), with colorized dots representing cells that expressed marker genes for each specified cluster, and gray dots representing cells that did not express these marker genes. Density plots depict the relative distribution of positively-identified cells in respective clusters over the mediolateral and dorsoventral extent of the VP. (**E**) Bar graphs indicating the density of detected cells belonging to individual clusters per mm^2^ over the anteroposterior axis of the VP.

Among the GABAergic subtypes (Fig 2A-C), the population marked by *Calcr* exhibited a bias of expression towards the medial extent of the VP, while *Sst*-positive neurons were relatively more concentrated in the most posterior extent of the VP (Fig. 2D). The two identified *Slc17a6*-positive clusters comport with publicly available MERFISH datasets; the SLC_MASP1 minority cluster is distinguished from the SLC_TCF7L2 majority cluster by exclusive expression of *Trp73* (Fig 3A-C); both populations exhibited the greatest relative expression in the medial antero-posterior levels of the VP (Fig 3D). Consistent with prior work, *Chat*- and *Cck*-positive cells were distributed sparsely throughout the VP, similar to adjacent basal forebrain structures (*30*, *31*). Overall, there was a lack of evident topographical organization of transcriptionally-defined VP cell types. Likewise, a separate analysis of dopaminoceptive neurons (defined by expression of the dopamine receptors *Drd1, Drd2,* and *Drd3*) revealed that all receptors spanned the extent of the VP (Fig. S3A), with *Drd1*- and *Drd2*-co-expressing nuclei and *Drd3*-positive nuclei expressed at relatively higher densities in the anterior VP (Fig S3B). Finally, immunohistochemistry corroborated the relative expression and general distribution of neuronal and non-neuronal classes determined by snRNA-seq in our mouse samples (Fig. S4).

### NHP atlases of ventral pallidum

The basal ganglia are highly conserved throughout vertebrates, yet the cellular composition of the VP remains largely unexplored in species other than rodents. As NHPs exhibit evolutionary similarity to humans, they serve as an optimal model organism for bridging the gaps between rodent and human studies and unraveling the underlying principles governing the transcriptomic profiles of basal ganglia neurons. We collected multiple VP samples (each hemisphere was collected separately) from individual rhesus macaque (n = 3; 1 male and 2 female) and baboon brains (n = 2 females) that were first perfused with ice cold saline and then cut into coronal slabs (2-4 mm thick) using custom 3D printed brain matrices. Landmarks on the ventral surface of the brain and the melanin-rich tissue appearing below the anterior commissure was used to guide dissection (Fig S5) based on standardized nonhuman primate atlases (*39*, *40*). We first conducted independent quality control and clustering analysis for each species (Table S1). The baboon dataset contains 57,724 high-quality nuclei with an average sequencing depth of 2,030.4 genes per nucleus (Fig. S6A) and the macaque dataset contains 49,542 high-quality nuclei with an average sequencing depth of 2,269.6 genes per nucleus (Fig. S6B). To identify transcriptionally distinct cell types, we first performed independent clustering of the snRNA-seq data from each species. In both macaque and baboon VP atlases, we identified six cell types including neurons (27.7% of total macaque nuclei, and 28.2% of total baboon nuclei), astrocytes (9.4% and 6%), microglia (11.8% and 15.4%), oligodendrocytes (41.8% and 37.3%), OPCs (8.7% and 12.9%), and vascular cells (0.6% and 0.2%) (Fig. 4A). These cell types exhibit high expression levels of previously reported canonical marker genes (Fig. 4B), which were largely conserved between the two NHPs as well as with mice. We also observed proportionally fewer neuronal nuclei in both NHP atlases compared to the mouse atlas, consistent with previous reports (*26*, *27*, *41–44*).

**Fig. 4.**
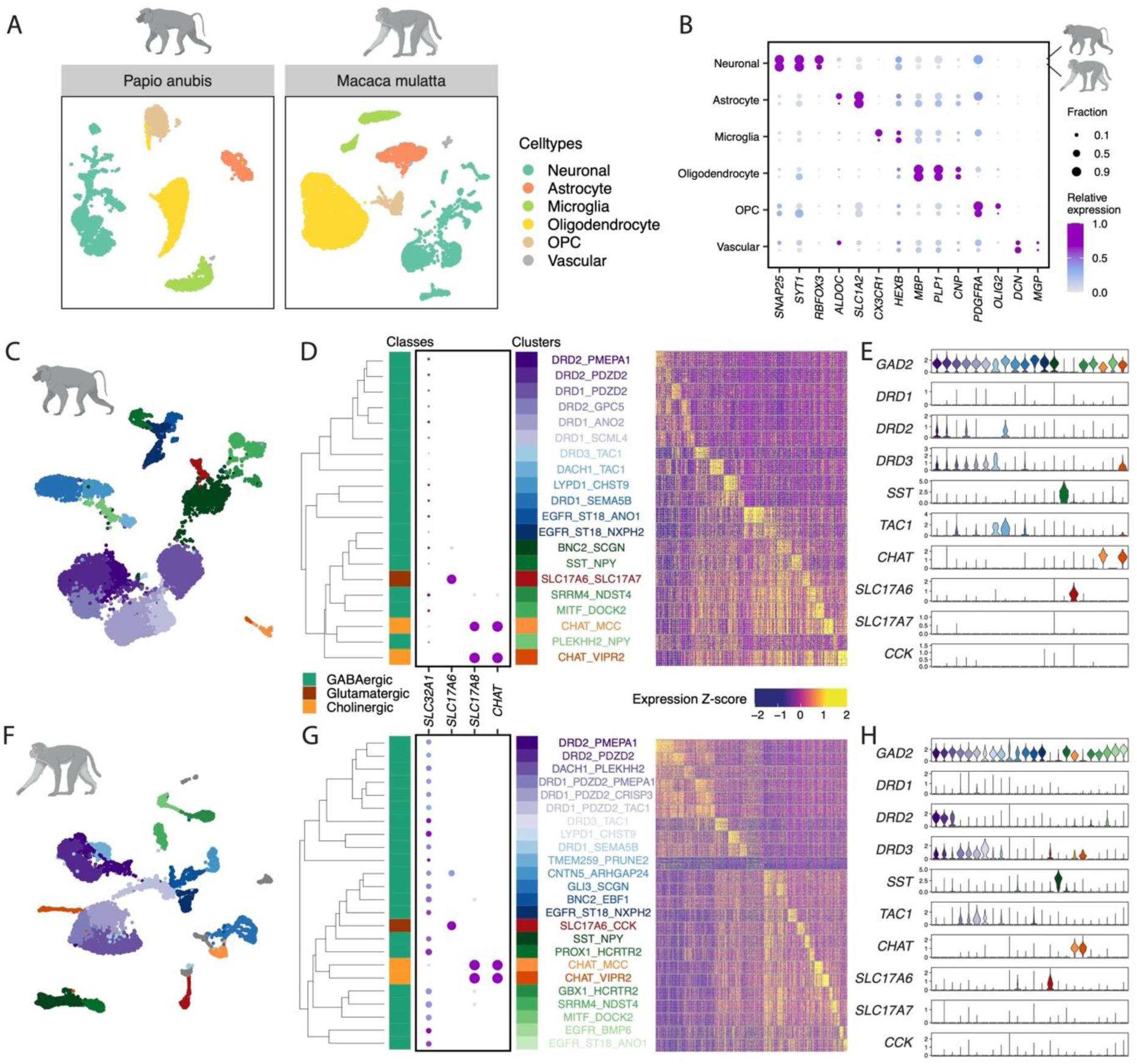
Atlases of the non-human primate ventral pallidum are revealed by snRNA-seq. (**A**) UMAP visualization of 10,000 downsampled nuclei each from baboon and macaque, colored by cell types. (**B**) Dot plot displaying the expression of select cell-type-specific marker genes in individual cell types from baboon and macaque. Dot size denotes the fraction of nuclei expressing a marker gene (>0 counts), and color denotes relative expression of a gene in each cell type (calculated as the mean expression of a gene relative to the highest mean expression of that gene across all cell types in the respective species). (**C** and **F**) UMAP of 10,000 downsampled neuronal nuclei from baboon (C) and macaque (F), colored by neuronal clusters. (**D** and **G**) Left: hierarchical clustering of neuronal clusters based on top 100 marker genes (sorted by Log2FC) per cluster in baboon (D) and macaque (G). Middle: Dot plot displaying the expression of select neurotransmitter genes for class annotation. See Fig 4B for legend. Right: Heatmap showing Z-scores of the expression of top 100 marker genes (sorted by Log2FC) per neuronal clusters in individual nuclei from baboon (D) and macaque (G). 50 nuclei per neuronal cluster are downsampled and displayed on the heatmap for each species. (**E** and **H**) Violin plot showing log-transformed expression of select marker genes in individual baboon (E) and macaque (H) neuronal clusters. Neuronal clusters are ordered according to the dendrogram in D and G, respectively.

Neuronal nuclei in each NHP species displayed high levels of transcriptional heterogeneity, a pattern similar to that found in our mouse VP atlas. To better understand the transcriptional heterogeneity of NHP VP neurons, we performed independent subclustering of the neuronal nuclei from each species. In baboon, we identified 20 neuronal clusters including 17 GABAergic clusters (97.3% of the neuronal nuclei) and three glutamatergic clusters, two of which are also cholinergic (Fig. 4C,D). In macaque, subclustering revealed 24 neuronal clusters including 21 GABAergic clusters (93.4% of the neuronal nuclei) and three glutamatergic clusters, two of which are cholinergic (Fig. 4F,G). Similar to what we found in the mouse, a small fraction of NHP neuronal nuclei co-express the GABAergic and glutamatergic neuronal markers. In macaque, 0.7% of neuronal nuclei co-express *SLC32A1* with one of the vesicular glutamate transporters. These nuclei were enriched in four clusters (Fig. S6K). In baboon, *SLC32A1* expression is low and only labels 1.8% of neuronal nuclei, compared to 18.4% in macaque and 30.9% in mouse. We, therefore, used *GAD1*, which labels 91.7% *SLC32A1*-expressing baboon nuclei (Fig. S6I), to estimate the population co-expressing GABAergic and glutamatergic neuronal markers. In baboon, 1.5% of neuronal nuclei co-express *GAD1* with one of the vesicular glutamate transporters and two clusters are highly enriched with the double-positive population (Fig. S6J).

In macaques and baboons, the clusters of GABAergic neurons in each atlas expressed high levels of *PDZD2* and were hierarchically organized based on transcriptional similarity, consistent with the observation in the mouse atlas. High expression of PDZD2 across multiple clusters of GABAergic neurons was consistent with what we found in the mouse atlas. But we did note a greater proportion of the total neuronal nuclei expressed PDZD2 in the baboon (58.8%) and macaque (50.2%) than in the mouse (32.9%) (Fig. S1C, S6G,H). Akin to the mouse atlas, we also identified clusters of glutamatergic and cholinergic neurons in the NHP VP. In both baboon and macaque, we identified two cholinergic clusters (CHAT_MCC and CHAT_VIPR2). These clusters also expressed *SLC17A8* (VGLUT3) indicating they were also glutamatergic neurons (Fig. S5G,H)(*34*). We also identified a distinct population of *SLC17A6* (VGLUT2)-positive neurons that co-localized with expression of CCK (cluster SLC17A6_SLC17A7 in the baboon, and SLC17A6_CCK in the macaque). While neurons of this cluster exclusively co-localized with *CCK* in the two NHP species, *SLC17A7* (VGLUT1) is uniquely expressed in the baboon (Fig. 4D,E, Fig. S6G). However, *SLC17A7* detection is low in the NHP atlases (less than 0.5% neuronal nuclei in each species, Fig. 4G,H) and future studies are needed to determine if the spatial localization of these different cell types is consistent across mammalian species.

Despite largely conserved marker genes that identify NHP and mouse VP neuronal clusters, there is a species-specific diversification of gene expression in VP neuronal subtypes. Macaque and baboon VP datasets revealed lower transcript detection of *DRD1*, *SLC17A7,* and *CCK*, and higher detection of *DRD3* relative to the mouse VP atlas. In addition, we observed higher expression of *TAC3*, the gene uniquely expressed in primate GABAergic interneurons (*44*), than we would have predicted based on the expression of the mouse homologue gene *Tac2. TAC3 is*, respectively, expressed in 6.3% and 8.6% of baboon and macaque neuronal nuclei, compared to *Tac2* expression in 3.6% of mouse neuronal nuclei. In both macaque and baboon VP atlases, *TAC3* is highly expressed in neuronal cluster EGFR_ST18_NXPH2 (Fig S6 G,H). Overall, when we examine each species independently, qualitative comparisons based on marker gene expression indicate that the VP transcriptional landscapes are largely shared across species. We next sought to harmonize these individual atlases to identify which cell types were conserved or unique to each species.

### Conservation of ventral pallidum neuronal populations across species

To take full advantage of the cross-species information and examine the correspondence of transcriptional cell types, we generated a harmonized atlas by integrating the neuronal nuclei from snRNA-seq data across all three species (*45*). Integration matched shared cell types across species as nuclei of the same major neuronal classes show extensive intermingling on the integrated UMAP compared to complete separation on the UMAP pre-integration (Fig. 5A, S7A). We next performed clustering of this harmonized neuronal atlas and generated 26 integrated clusters (Fig. S7B).

**Fig. 5.**
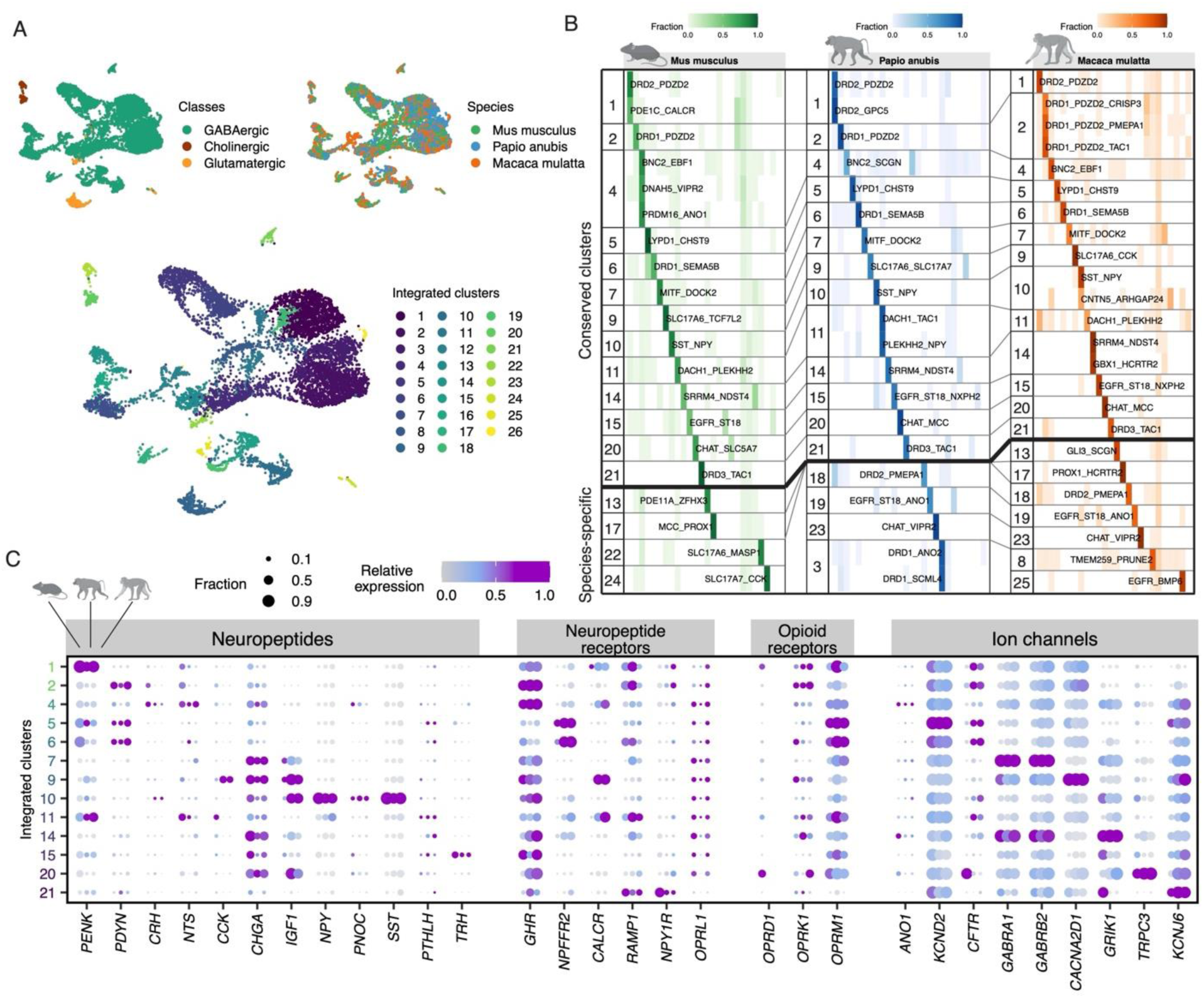
Harmonization of mouse and non-human primate neurons reveals conservation of transcriptional cell types across species. (**A**) UMAPs of the harmonized neuronal atlas of VP. 3,000 neuronal nuclei per species are downsampled from mouse, baboon, and macaque. Nuclei are colored by neuronal cell types (top left), species (top right), and integrated clusters from the harmonized atlas (bottom). (**B**) Heatmap showing the correspondence of neuronal clusters across species. Fraction of nuclei in each neuronal cluster defined by independent analysis for each species that mapped to each integrated cluster identified in the harmonized atlas is calculated. Integrated clusters consisting of nuclei that made up over 30% of the nuclei from at least one neuronal cluster per species are considered conserved clusters. All other integrated clusters are identified as species-specific. (**C**) Dot plot displaying the expression of select gene panels in individual conserved integrated clusters from mouse, baboon, and macaque. Dot size denotes the fraction of nuclei expressing a marker gene (>0 counts), and color denotes relative expression of a gene in each cell type (calculated as the mean expression of a gene relative to the highest mean expression of that gene across all cell types in the respective species). Opioid receptor *OPRD1* or its homologues are not annotated in the two NHP reference genomes.

To identify neuronal cell types that are evolutionarily conserved, we first examined the integrated clusters that showed a high degree of correspondence to individual neuronal clusters from each of the species characterized by independent analysis. We identified 13 clusters in the integrated atlas that each had at least one corresponding neuronal cluster in every species (Integrated clusters 1, 2, 4, 5, 6, 7, 9, 10, 11, 14, 15, 20 and 21, Fig. 5B, S7C, Methods). Compared to all other neuronal clusters identified based on independent analysis, neuronal clusters of different species within a conserved cluster exhibit significantly higher transcriptional similarity with respect to the expression of conserved marker genes (Pearson’s r = 0.87 ± 0.12 among neuronal clusters in the same integrated clusters, compared to Pearson’s r = 0.62 ± 0.11 among neuronal clusters not in the same integrated clusters, p = 3.94e-63, two-sided student t-test; Fig. S6D). Many of these conserved clusters are identified by genes encoding neuropeptides or neuropeptide receptors, i.e. DRD2_PDZD2, DRD2_GPC5, DRD1_PDZD2, DRD1_SEMA5B, SST_NPY, CHAT, DRD3_TAC1 (Fig. 5B, Table S5), and the expression of these functionally relevant genes is similar in nuclei within the conserved clusters in all three species (Fig. 5C). This indicates that these 13 integrated clusters represent evolutionary conserved neuronal populations, constituting 57.6%, 61.7%, and 60.9% of neuronal nuclei in mice, baboons, and macaques, respectively.

Among the conserved integrated clusters, 11 were GABAergic. For example, integrated cluster 1 is conserved, consisting of nuclei with 38.3%, 39.8%, and 21.9% from mouse, baboon, and macaque, respectively. Mouse nuclei of this integrated cluster constitute 85.4% of the nuclei in mouse cluster DRD2_PDZD2 and 59.6% of the nuclei in mouse cluster PDE1C_CALCR. Baboon nuclei of this integrated cluster constitute 89% of the nuclei in baboon cluster DRD2_PDZD2 and 97.7% of the nuclei in baboon cluster DRD2_GPC5. Macaque nuclei of this integrated cluster constitute 86.7% of the nuclei in macaque cluster DRD2_PDZD2 (Table S5). These data suggest that integrated cluster 1 represents a conserved neuronal population across three species and that the neuronal clusters, DRD2_PDZD2 and PDE1C_CALCR in mouse, DRD2_PDZD2 and DRD2_GPC5 in baboon, and DRD2_PDZD2 in macaque, display high transcriptional correspondence. Nuclei of integrated cluster 1 from different species exhibit conserved transcriptional profile of their marker gene expression (Fig. S7D) as well as functionally relevant gene panels (Fig. 5C).

The harmonization also revealed two conserved clusters of glutamatergic neurons. In the conserved integrated cluster 9, the mouse cluster SLC17A6_TCF7L2 exhibited high transcriptional correspondence to SLC17A6_SCL17A7 in baboon and SLC17A6_CCK in macaque. Integrated cluster 20 represented a conserved cholinergic (also glutamatergic based on the expression of *Slc17a8*/*SLC17A8*) neuronal population. Mouse cholinergic neurons formed a single cluster, CHAT_SLC5A7, and 73.2% of the nuclei from this mouse cholinergic cluster exhibited high transcriptional correspondence with the CHAT_MCC clusters in NHPs.

### Divergent ventral pallidum neuronal populations

Despite being largely conserved, VP neurons also exhibit several species-specific characteristics. We identified 10 integrated clusters that were not conserved across all three species (Integrated clusters 3, 8, 13, 17, 18, 19, 22, 23, 24, 25, Fig 5B). To investigate the evolutionarily divergent VP features, we first identified three integrated clusters that are conserved between two NHP species with no homologue in the mouse (Integrated clusters 18, 19, and 23). Integrated cluster 18 was conserved across NHPs and marked by expression of *DRD2* and *PMEPA1*. *PMEPA1* is an androgen responsive gene that has been characterized for its essential and reciprocal regulation of androgen receptor (AR) signaling (*46*). Additional genes that differentiated the DRD2_PMEMPA1 cluster with other neuronal clusters in both NHP species included *ACVR1*, *SORCS3, FOXN3, FGF13,* and *MOXD1*, which are critical for axonal remodeling and synaptic regulation (Fig. S7E). This raises the possibility that this NHP conserved cluster represents a unique transcriptional cell state reflecting the increased neuronal plasticity of NHPs. Integrated cluster 19 is enriched of EGFR_ST18_ANO1 neurons from both baboon and macaque. In both NHPs, EGFR_ST18_ANO1 neurons highly expressed *TAC3* and *VIP*, as well as *NHX2-1* and *LHX6* (Fig S7G). In mouse, *Tac2* (*TAC3* homologue) and *Vip* are associated with interneurons of a caudal ganglionic eminence (CGE) origin (*47*), and *Nhx2-1* and *Lhx6* are transcription factors expressed in interneurons of a medial ganglionic eminence (MGE) origin. The gene expression profile of EGFR_ST18_ANO1 neurons is consistent with a novel primate-specific MGE-derived interneuron population that also expresses CGE-associated neuropeptides, previously described in marmoset and human striatum (*44*). Moreover, we identified a second distinct NHP cholinergic cluster (integrated cluster 23), corresponding to CHAT_VIPR2 in both macaques and baboons. This NHP neuronal population corresponds less strongly to the mouse cholinergic neurons compared to the conserved CHAT_MCC population. The NHP CHAT_VIPR2 cluster is marked by expression of the vasopressin receptor *VIPR2*, raising the possibility that these populations are differently subjected to excitation by vasopressin from exogenous sources or by local release from VP neurons (Fig. S7F). *Vipr2* is not notably expressed in mouse cholinergic neurons.

While some glutamatergic neuronal clusters are transcriptionally conserved as described above, glutamatergic neurons overall represented a higher proportion of VP neurons in mice (9.9% of mouse neuronal nuclei, Fig 1E), relative to either primate species (2.7% of baboon neuronal nuclei and 6.6% of macaque neuronal nuclei, Fig 4D,G). In addition, some glutamatergic populations exhibited divergence across species. The minor glutamatergic clusters in mouse, SLC17A6_MASP1 (integrated cluster 22) and SLC17A7_CCK (integrated cluster 24), had no transcriptional homologue in primate species, suggesting these clusters diverge between mouse and NHP.

## Discussion

The basal ganglia are highly conserved throughout vertebrates, and our results underscore this conservation of the VP across rodents and primates at the transcriptional level. The VP is classically considered a GABAergic nucleus with cholinergic neurons which comprise part of the basal forebrain cholinergic system (*14*, *48–54*) while more contemporary work has identified that glutamatergic neurons comprise a distinct subpopulation (*6*, *7*). Our results confirm that the division of VP neurons into these broader neurochemical classes is valid and holds across mammalian species. Each of these neurochemical classes is comprised of distinct transcriptional subtypes, which are also largely conserved across mammalian species. Intriguingly, the proportion of neurons represented in each of these subclasses diverged between species; while putative GABAergic neurons comprised the majority of VP neurons in all species, primates exhibited a smaller proportion of glutamatergic neurons and a larger proportion of cholinergic cells relative to mice. VP cholinergic neurons project to either the frontal cortex or amygdala (*55*), future work will establish whether the two distinct cholinergic clusters in NHPs map onto these projection targets. Alternatively, since only one cholinergic cluster had a homologous cluster in mice, our results could indicate evolutionary drift in the neuronal subtypes essential for the specialized functional role in the NHP. This diversification in the NHP cell types could also be a product of the VP expansion or possible integration of new sensory signals in the VP (*56–64*).

We identified 20 neuronal subtypes in mouse VP, marked by enrichment of neuropeptides, neurotransmitter receptors, and transcription factors. While foundational work has leveraged Cre driver lines (i.e. *Slc32a1*, *Slc17a6*, *Pvalb*, *Drd3*) to dissect VP circuit function, our results argue that these strategies do not correspond to molecularly defined VP subtypes. One framework for interpreting the heterogeneity of the VP is that VP cell types represent a combination of neurons from adjacent structures, including the ventral striatum and extended amygdala (*65*). Our results corroborate this interpretation at the transcriptional level. Specifically, we identified VP populations in mice that share marker genes for accumbal medium sized spiny neurons (*PDZD2* co-expressed with either *DRD1* or *DRD2* (*32*, *33*, *66*), clusters with transcriptional homologues in the extended amygdala (SLC17A7_CCK, DNAH5_VIPR2)(*30*, *67*, *68*), striatal interneurons expressing *Tac3* (EGFR_ST18_ANO1) and glutamatergic neurons with transcriptional homology to those expressed in BNST (SLC17A6_MASP1) (*28*, *32*, *33*), and GABAergic Chst9-positive neurons (LYPD1_CHST9) that are highly enriched in mu opioid receptors as previously reported in the nucleus accumbens (*69*). Recent reports have corroborated this interpretation, finding that transcriptionally defined cell types are expressed in multiple forebrain structures that share borders with the VP (26, 33, 34).

A potential limitation of our study is that the VP is elongated and irregularly shaped, and our tissue punches likely did not capture the entire structure in all samples and species. Our histological analysis confirmed that the rostral and posterior subcommissural VP territories were represented in our mouse and NHP samples. Therefore, it is possible that cell types that may be highly expressed in the most anterior sections of the VP are underrepresented in our samples, due to difficulty distinguishing the sparse regions of VP from the intervening nucleus accumbens. The interpeduncular nucleus (IPAC) (*70*), or fundus of the striatum (*33*) abuts the VP laterally, and it is possible that in our most posterior punches, neurons from this structure were included. Our spatial validation confirms that this area is highly enriched in *Pdzd2* and *Slc17a7*; however, the small proportions of these cell types in our sequencing dataset, and orthogonal confirmation of the expression of *Pdzd2* and *Slc17a7* sparsely throughout the VP argue against excessive contamination from adjacent structures.

Our spatial validation also revealed an absence of clear topographical organization of neuronal populations marked by GABA, glutamate, and acetylcholine, with a lack of discrete boundaries between cell types nor continuous gradients of gene expression. However, we uncovered some evidence for spatial organization of the specific sub clusters of GABAergic and glutamatergic clusters. Prior pharmacology and functional studies have demonstrated functional anatomical subdivisions of VP. For example, infusion of opioids into the posterior VP enhances hedonic reactions to palatable stimuli, whereas infusion into the anterior VP suppresses these same reactions. Relatedly, GABA-mediated inactivation of the VP promotes feeding, with effects more pronounced when medial VP is targeted (*71*). Our results suggest that differences in topography of transcriptionally-defined and dopaminoceptive cell types across the anterior-posterior gradient within the VP may contribute to these functional differences.

The VP is composed of non-overlapping populations of anatomically-defined projection neurons to the nucleus accumbens, lateral habenula, thalamus, cortex, and midbrain which have been causally linked to distinct dimensions of reward processing(*9*, *72–77*). It will be critical to know the anatomical identity of these transcriptionally-defined populations to understand how these cell types modulate behavior. Moreover, transcriptomes are not static, responding to the local environment even in terminally differentiated, post-mitotic neurons. Given interest in the VP as a target for depression(*9*, *78*), obesity(*11*, *79*) and substance use disorders(*2*, *80*, *81*), future work necessitates understanding how experience and environmental stressors alter the transcriptional profile of these VP cell types, and ultimately how these transcriptional changes alter neuronal and circuit function.

## Methods

### Subjects

#### Mouse

C57Bl6J mice (aged 10-15 weeks) were used for snRNA-sequencing (n=10 males, 10 females), in situ hybridization (2M, 2F), and immunohistochemistry (2M, 2F) experiments. Mice were housed in groups of 2-5 with ad libitum access to standard rodent lab chow and water, on a 12h light cycle (light onset 7 am, offset 7 pm). All procedures were approved by the institutional animal care and use committee at Washington University School of Medicine (WashU). Study approval numbers: 21-0299 (WashU)

#### Primates

2 male olive baboons (3.62 and 4.32 years old), 1 male rhesus macaque (10.7 years old), and 3 female rhesus macaques (5.87, 6.66, and 17.67 years old). All animals were pair housed on a 12-hour on/12-hour off lighting schedule with ad libitum access to food and water. Animals were fed standard primate chow twice daily and provided fruit and vegetable enrichment daily. Macaques were observed by trained veterinary technicians daily in their home cages. The Institutional Animal Care and Use Committee and the Institutional Biosafety Committee at the ONPRC and Oregon Health and Science University (OHSU) approved all experimental procedures. All of the guidelines specified in the National Institutes of Health Guide for the Care and Use of Laboratory Animals (National Research Council, 2011) were strictly followed. Study approval numbers: IP00002202 (OHSU).

### Mouse Tissue Collection for snRNA-seq

To collect mouse VP tissues for sequencing, animals were anesthetized with ketamine cocktail, and perfused with ice-cold NMDG-based cutting solution (NMDG 93 mM, KCl 2.5 mM, NaH_2_PO_4_ 1.25 mM, NaHCO_3_ 30 mM, HEPES 20 mM, Glucose 25 mM, Ascorbic acid 5 mM, Thiourea 2 mM, Sodium Pyruvate 3 mM, MgSO_4_ 10 mM, CaCl_2_ 0.5 mM, N-acetylcysteine 12 mM; pH adjusted to 7.3 with 12N HCl, and bubbled with 95% O_2_ and 5% CO_2_). Brain was dissected submerged in ice-cold NMDG-based cutting solution and sliced coronally into sections with 400 µm thickness using a Compresstome (Precisionary, VF-210-0Z). VP regions were micro-dissected under a microscope (Leica S9i) in a sterile petri dish treated with RNAse-X on dry ice, using a reusable 0.75 mm biopsy punch (WPI 504638). For each sequencing sample, bilateral VP tissues from five mice were punched and pooled into a nuclease-free centrifuge tube placed on dry ice. Given the irregularity and lack of clear boundaries between the VP and the nucleus accumbens (anterior VP) and substantia innominata (posterior VP), samples were collected between + 0.6 mm and −0.2 mm from Bregma, immediately ventral to the anterior commissure. Samples were stored at −80 °C before nuclei isolation.

### Primate Tissue Collection for snRNA-seq

Freshly-dissected, unfixed, and flash-frozen VP tissue punches were acquired from two non-human primate species. Necropsies and tissue collections were performed as previously described(*82–84*). Animals were sedated with ketamine (10 mg/kg) and then deeply anesthetized with sodium pentobarbital followed by exsanguination. The brain and spinal cord were perfused through the ascending carotid artery with 1 L of 0.9% ice-cold saline. The brain was then removed from the skull (< 30 minutes post-mortem), deposited into an ice-cold bath of saline for transport, and placed into an ice-cold, steel brain matrix (Electron Microscopy Sciences). A custom 3-D printed brain matrix was used to section the brains. Each brain was positioned in the brain matrix with the ventral surface facing up. The anterior medial temporal sulcus, posterior to the temporal pole, and anterior to the rhinal sulcus was identified on the ventral surface of the temporal lobe. A carbon steel knife blade (Thomas Scientific) was inserted into the slot in brain matrix that was most closely aligned and orthogonal to the beginning of the anterior medial temporal sulcus. Additional knife blades were inserted anterior and posterior to the first knife blade inserted in 2 mm increments. The resulting 2 mm bilateral brain slabs were then removed from the brain matrix and laid out flat in sterile petri dishes treated with RNase-X and stored at −80°C. The petri dishes rested on an aluminum plate secured to a chamber filled with dry ice. The plate temperature was maintained at −30°C to −15°C for up to two hours while tissue punches were acquired and flash frozen.

Brain slabs in which the VP was clearly visible below the anterior commissure on both sides of the slab were identified and 1.0-2.5 mm in diameter tissue punches (1-3 per site) were taken through the full width of the slab (Fig S5). Tissue punches from each hemisphere were checked to ensure they did not contain white matter and then inserted into DNase- and RNase-free 1.5 mL microcentrifuge tubes that were inserted into powderized dry ice. Ethanol was poured onto the dry ice to flash freeze the tissue punches. All tissue punches were acquired within 90 minutes postmortem. The punched brain slabs were then post fixed in 4% paraformaldehyde for 48 hours, cryoprotected in 30% sucrose, and then sectioned in 40 µm sections for histological analyses.

### Single-nuclei isolation

Nuclear extraction for both mice and primates was performed with density gradient centrifugation according to the protocol described previously with modifications to scale down the reaction volume (*85*). Samples were moved from −80 °C and thawed on ice for 2 minutes. Tissues were transferred to a dounce homogenizer (Millipore Sigma, D8938-1SET) in homogenization buffer (0.25 M sucrose, 25 mM KCl, 5 mM MgCl_2_, 10 mM Tris-HCl, pH 8.0, 5 µg/mL actinomycin, 1% BSA, and 0.08 U/µl RNase inhibitor, 0.01% NP40) on ice. Samples were homogenized for 10 strokes with the loose pestle in a total volume of 1 mL, followed by 10 additional strokes with the tight pestle. The tissue homogenate was then passed through a 50 µm filter and diluted 1:1 with working solution (50% iodixanol, 25 mM KCl, 5 mM MgCl_2_, and 10 mM Tris-HCl, pH 8.0). Nuclei were layered onto an iodixanol gradient after homogenization and ultracentrifuged as described previously. After ultracentrifugation, nuclei were collected between the 30 and 40% iodixanol layers and diluted with resuspension buffer (1xPBS with 1% BSA, and 0.08 U/µl RNase inhibitor). Nuclei were centrifuged at 500 g for 10 min at 4°C and resuspended in resuspension buffer with 5 ng/µl of 7-AAD. FACS was carried out to further remove cellular debris. 7-AAD+ events were collected using a 100 µm nozzle on a BD FACSAria III instrument into a 1.5 mL microcentrifuge tube.

### snRNA-seq

Nuclei were counted using a hemocytometer and diluted to the concentration that targets 10,000 nuclei recovery. Nuclei were processed according to the manufacturer’s manual of 10X Genomics Chromium Single Cell Gene Expression 3’ V3.1 Assay. Libraries were sequenced on a NovaSeq6000 with 150 cycles each for Read1 and Read2, targeting 50,000 paired reads/nucleus. Raw sequencing data from individual libraries were processed and mapped to the reference genome (mouse: mm10, baboon: Panubis1.0, and macaque: Mmul_10) and the gene-cell counts matrices from all snRNA-seq libraries of the same species were concatenated using 10X Genomics cellranger-v6.1.2 or cellranger-v7.0.1.

#### Quality control and clustering of snRNA-seq

The analysis was carried out using R package Seurat (V4.3.0) (*86*) with data derived from different species analyzed separately. Nuclei with at least 500 unique genes, less than 15,000 unique transcript counts, and fewer than 2% for mouse and 5% for baboon and macaque of the counts from mitochondrial genes were included in the analysis. Raw counts were scaled to 10,000 transcripts per nucleus using NormalizeData() function to control the sequencing depth between nuclei. Counts were centered and scaled for each gene using ScaleData() function. Highly variable genes were identified using FindVariableFeatures() and the top 20 principal components (PCs) were retrieved with RunPCA() using default parameters. For dimension reduction and visualization, Uniform Manifold Approximation and Projection (UMAP) coordinates were calculated based on the top 20 PCs using RunUMAP(). Nuclei clustering was performed using FindClusters() based on the variable features from the top 20 principal components, with resolution of 0.6, and the marker genes for each cluster were identified using FindAllMarkers() comparing nuclei in one cluster to all other nuclei. Clusters enriched of low-quality nuclei or doublets were identified if they meet any of the following criteria: 1). Assigned to a cluster with no significantly enriched marker genes (FDR < 0.05, log2FC > 1); 2). Assigned to a cluster in which five or more mitochondrial genes were identified among the top 20 marker genes (sorted by avg_log2FC). The remaining clusters were either assigned as neuronal if they expressed high level of neuronal marker genes (*Rbfox3*, *Snap25*, and *Syt1*), or as non-neuronal if they exhibited high expression level of any of the non-neuronal marker genes (*Sparc*, *Mbp*, or *Apoe*). Nuclei from neuronal clusters and non-neuronal clusters were sub-clustered separately, and the clusters enriched with low-quality nuclei or doublets were identified again as described previously.

#### Mouse libraries

For the mouse dataset, each biological sample was prepared by pooling bilateral VP tissues from five mice. Four biological samples (two female samples and two male samples) were loaded into five sequencing libraries (with one sample loaded into two libraires), generating a total of 44,518 nuclei that passed initial quality control cutoffs (Table S1). Among them, 6,388 nuclei were identified as doublets or low-quality and thus excluded from the dataset, yielding a final dataset containing 38,130 nuclei (24,870 neuronal nuclei and 13,260 non-neuronal nuclei).

#### Baboon libraries

For the baboon dataset, VP punches from each hemisphere of two animal subjects were separately collected, generating four biological samples. They were loaded into six sequencing libraries (with one sample loaded into three libraires), generating a total of 60,209 nuclei that passed initial quality control cutoffs (Table S1). Among them, 2,485 nuclei were identified as doublets or low-quality and thus excluded from the dataset, yielding a final dataset containing 57,724 nuclei (16,293 neuronal nuclei and 41,431 non-neuronal nuclei). While baboon nuclei from individual libraries clustered together on the UMAP indicating a low batch effect across libraries (Fig. S6C), snRNA-seq libraries of one animal identified much fewer neuronal nuclei compared to those of another animal (5.6 ± 0.8% compared to 36.1 ± 11.8%, mean ± standard deviation, Fig. S6E). This difference in neuronal nuclei enrichment was likely due to variations from individual animals (e.g. age or other individual factors) and/or tissue procurement process.

#### Macaque libraries

For the macaque dataset, VP punches from each hemisphere of three animal subjects were separately collected, generating four biological samples (one hemisphere of two subjects and both hemispheres of one subject). They were loaded into six sequencing libraries (with one sample loaded into three libraires), generating a total of 49,802 nuclei that passed initial quality control cutoffs (Table S1). During the initial clustering, individual macaque snRNA-seq libraries showed similar cell type compositions (Fig. S6F) and clustered together on the UMAP space (Fig. 65D), with the exception that one snRNA-seq library exhibited mild transcriptional variation compared to other libraries (Fig. S6D, Table S4). This altered gene expression to this library, likely due to technical variation in the tissue extraction and procurement process. To correct for the batch effect, the gene expression matrices from individual libraries were normalized using FindIntegrationAnchors() and integrated using IntegrateData(), both with the normalization method set as ‘SCT’. Clustering was carried out as described above. Among them, 260 nuclei were identified as doublets or low-quality and thus excluded from the dataset, yielding a final dataset containing 49,542 nuclei (13,696 neuronal nuclei and 35,846 non-neuronal nuclei).

### Cluster Annotation

Clusters were annotated and named based on the expression of canonical markers for each cell type: neurons (*Rbfox3, Snap25, Syt1* mouse; *RBFOX3, SNAP25, SYT1* primate), astrocytes (*Aldoc*, *Aqp4*, *Slc1a2* mouse; *ALDOC*, *AQP4*, *SLC1A2* primate), choroid epithelial cells (*Ttr*, *Folr1*, *Clic6* mouse; *TTR*, *FOLR1*, *CLIC6* primate), macrophages (*Pf4, Cd74, Mrc1* mouse; *PF4, CD74, MRC1* primate), microglial cells (*Tmem119, Cx3cr1, Hexb* mouse; *TMEM119, CX3CR1, HEXB* primate), oligodendrocytes (*Mbp, Plp1, Cnp* mouse; *MBP, PLP1, CNP primate*), OPC (*Pdgfra, Olig1, Olig2* mouse; *PDGFRA, OLIG1, OLIG2* primate), and vascular cells (*Dcn, Mgp, Cped1* mouse, *DCN, MGP, CPED1* primate).

To identify neuronal cell types, nuclei in the neuronal clusters were extracted to be re-clustered. Initial independent clustering of each species resulted in 20 clusters in mice, 20 clusters in baboon, and 24 clusters in macaque. Each cluster was assigned a neurochemical class based on the canonical marker gene that is most predominantly expressed in each cluster. First, a cluster is classified ‘cholinergic’ if the average expression of *Chat* (mouse, or *CHAT* primate) across all nuclei in the cluster is greater than its average expression plus two standard deviations across all neuronal nuclei of the given species. Next, the remaining clusters are classified as ‘glutamatergic’ if the average expression of either *Slc17a6*, *Slc17a7*, or *Slc17a8* (mouse, or *SLC17A6* or *SLC17A8* primate, *SLC17A7* expression in primate is low and thus not used for classification) is greater than the average expression of the corresponding gene plus two standard deviations across all neuronal nuclei of the given species, respectively. Finally, the remaining clusters are classified as ‘GABAergic’ if the average expression of *Slc32a1* (mouse, or *SLC32A1* primate) is greater than the 70% percentile of its expression across all neuronal nuclei of the given species. Clusters expressing multiple neurotransmitters were discussed in the results section. Clusters were further annotated and named based on differentially expressed marker genes conserved across species. Mouse cluster names are presented in uppercase to be consistent with primate names.

### Cross-Species Analysis

To harmonize VP neurons across species, neuronal nuclei from each species were selected. Mouse genes were converted to primate genes by uppercasing the gene symbols. We note that with this strategy, a possible limitation is that orthologous genes may have different symbols in different species, which may lead us to underestimate transcriptional similarity between clusters. Baboon, macaque, and mouse datasets were jointly clustered using Seurat V4 as described above (Fig. S4A). To facilitate accurate comparative analysis across species, nuclei of different species were integrated (Fig. 4A). Top 1000 variable genes in each species were used to generate an integrative expression matrix, and integration anchors were identified by running FindIntegrationAnchors(). Data of different species were then integrated using IntegrateData(). The top 20 PCs were used for clustering and UMAP visualization. Integrated clustering yielded 26 clusters. To establish correspondence between cross-species integrated clusters and mouse, macaque, and baboon clusters (Fig. 4B), which were generated independently by the clustering approach described above, we calculated the fraction of nuclei in each neuronal cluster defined by independent analysis for each species that mapped to each integrated cluster. Integrated clusters consisting of nuclei that made up over 40% of the nuclei from at least one neuronal cluster per species and at least 30 nuclei per species are considered conserved clusters. Integrated clusters 1, 2, 4, 5, 6, 7, 9, 10, 11, 14, 15, 20, and 21 were classified as conserved clusters. Remaining integrated clusters consisting of nuclei that made up over 40% of the nuclei from at least one neuronal cluster and at least 30 nuclei from at least one species are considered species-specific clusters. Integrated clusters 13 and 17 correspond to mouse and macaque neuronal clusters. Integrated clusters 22 and 24 correspond to only mouse neuronal clusters. Integrated clusters 18, 19, and 23 correspond to baboon and macaque neuronal clusters. Integrated cluster 3 only corresponds to baboon neuronal clusters, and integrated clusters 8 and 25 correspond to only macaque neuronal clusters. Remaining integrated clusters (12, 16, and 26) showed no strong correspondence to any neuronal clusters.

To assess the transcriptional similarity of cell types across different species (Fig. S6D), expression values of conserved marker genes (Log2FC>0.5, FDR<0.05 in all species) in all nuclei of each neuronal cluster identified by independent analyses were averaged for each gene using AverageExpression() and then pairwise Pearson’s correlation coefficients between any pairs of clusters in each dataset were computed.

### Differential Expression Analysis

Differential expression analysis was done using R package Seurat. Marker genes for each cell type were identified using FindAllMarkers() comparing nuclei in one cell type to all other nuclei. Marker genes for each neuronal cluster were identified using FindAllMarkers() comparing nuclei in one neuronal cluster to all other neuronal nuclei.

Differential expression analysis on sex-specific gene expression was performed on pseudobulk data to control technical variations across biological samples of the same sex. Specifically, pseudobulk counts for each neuronal cluster or non-neuronal cell type were generated by aggregating counts from nuclei of the same biological sample using AggregateExpression() in Seurat. Differential expression analysis was done using DESeq2 V1.44.0 in R by comparing counts of biological samples between sex (n = 2 each sex). Genes with FDR < 0.05 were reported.

### Gene Ontology (GO) Analysis

GO analysis was performed using topGO (version: 2.40.0) in R. Conserved marker genes (log2FC > 0.5, FDR < 0.01 in all examined species) were used as the input gene list. For comparison, the background gene list included all genes with average expression > 0.5 in the respective nuclei being analyzed. Genes were annotated for their biological process and associated gene ontology terms. Enrichment is defined as the number of annotated genes observed in the input list divided by the number of annotated genes expected from the background list. GO terms with at least 10 annotated genes and enrichment p-value < 0.05 were returned except otherwise specified.

### Fluorescent *In Situ* Hybridization (HiPlex FISH)

#### Sample preparation

Mice were anesthetized with isoflurane, sacrificed, and brains were rapidly dissected then flash frozen in isopentane at −80°C. Flash-frozen brains were hemisected and sectioned at a thickness of 10 µm on a cryostat (Leica CM1850; Leica Biosystems, Deer Park, USA). Sections spanning the rostrocaudal extent of the VP were collected on Superfrost Plus microscope slides (Fisher Scientific, Pittsburgh, USA). Slides were stored at −80°C until further use. HiPlex probes for *Gad2, Drd1, Drd2, Calcr, Bnc2, Chst9, Lypd1, Sst, Tac1*, *Ano1, Prox1, Chat, Cck, Slc17a6, Trp73, Sox6, and Vip* mRNA were hybridized and fluorescently labeled using the RNAScope HiPlex12 Reagent Kit v2 (488, 550,650) Assay (Advanced Cell Diagnostics, Newark, USA) according to manufacturer’s instructions. Multiplex probes for *Drd3*, *Slc17a7*, and *Pdzd2* mRNA were hybridized and fluorescently labeled using the RNAScope Multiplex Fluorescent Reagent Kit v2 (520, 570, 650) Assay (Advanced Cell Diagnostics, Newark, USA) according to manufacturer’s instructions. DAPI was used as a counterstain and slides were coverslipped using Gold Prolong antifade mounting medium (Invitrogen, Eugene, OR).

#### Signal detection and analysis

Tilescans and z-stacks encompassing the entire VP were acquired using a confocal microscope (CSU-W1 SoRa spinning disk confocal; Nikon Instruments, Melville, USA) under the 20X objective. Channels collected across rounds of imaging were aligned in ACDBio HiPlex Registration software using the DAPI signal. Tiff files were then analyzed in QuPath (v0.5.1). Using the ‘Cell detection’ function, individual DAPI nuclei were segmented with a 2 µm expansion perimeter. The threshold detection level, sigma parameter, and minimum area were individually determined for each section. The ‘Subcellular detection’ function was then used and individual threshold levels for each channel was determined to obtain the approximate number of puncta for each probe. All detections were then exported and histograms showing the number of subcellular detections for each probe were generated per section. This allowed us to determine the number of detections required to classify a cell as positive for each marker. Each cell was then given an identity based on the probes that were found to be positively expressed, and cluster membership for each cell was assigned based on combinatorial co-expression of probes. Nuclear coordinates and corresponding cell identity were then mapped and aligning to anatomical landmarks of the VP according to the atlas of Paxinos and Watson ((*70*); R version 4.3.3).

### Immunohistochemistry

#### Sample preparation

Mice were anesthetized with isoflurane, PFA-perfused, and sacrificed. Brains were dissected and stored in 30% sucrose cryoprotectant at 4°C until brains were fully submerged. 30 µm sections through the rostrocaudal extent of the VP were collected using a cryostat (Leica CM1850; Leica Biosystems, Deer Park, USA) and then hemisected. The sections were placed in blocking solution (94.5% PBS, 5% Normal Donkey Serum, 0.1% Triton X) for one hour. Primary antibodies were allowed to incubate on an orbital shaker overnight at 4°C. Antibodies were diluted with blocking solution at the following concentrations: NeuN (ABN90, Sigma-Aldrich) 1:3000, Iba1 (234 009, Synaptic Systems Company) 1:1000, Sox9 (ab185966, Abcam) 1:500. Following primary antibody incubation, we performed three 10-minute washes in PBS, using fresh PBS for each wash. The following secondary antibodies were then applied and incubated for 2 hours at room temperature on an orbital shaker: Goat anti-guinea pig Alexa Fluor 647 (A-21450, ThermoFisher) 1:1000, goat-anti-chicken Alexa Fluor 488 (A-11039, ThermoFisher) 1:1000, donkey anti-rabbit Alexa Fluor 594 (A-21207, ThermoFisher) 1:1000. Sections were then mounted onto Superfrost Plus microscope slides (Fisher Scientific, Pittsburgh, USA) and coverslipped using Fluoromount-G, with DAPI (Invitrogen, USA).

#### Image acquisition and analysis

Images along the anterior-posterior extent of the VP were imaged on a Zeiss AxioImager.Z2 (Apotome) at 20x. ROIs were captured as 600,000 µm^3^ (50000µm^2^, 12 µm z-stacks) for quantification of NeuN, Iba1, and Sox9. Supervised pixel and object classification was performed using Ilastik machine-learning software (v1.4, (*87*)). Pixel classification was done by drawing annotative lines on cells, across the diameter of the cells, and on the background. After thresholding and filtering (hysteresis thresholding for NeuN and simple thresholding for Iba1 and Sox9), extracted objects were annotated as cells or backgrounds. We validated the Ilastik model by randomly selecting 10 ROIs and corroborating counts generated manually and in Ilastik. Images were then batch processed using the pixel and object classification models before exporting counts for NeuN (neurons), Iba1 (microglia), and Sox9 (astrocytes).

## Acknowledgments

We thank all members of the Tissue Distribution Program administered through Division of Comparative Medicine Pathology Services Unit at the Oregon National Primate Research Center for their assistance in acquiring nonhuman primate brain tissue. We thank Flow Cytometry & Fluorescence Activated Cell Sorting Core in the Department of Pathology and Immunology at Washington University School of Medicine for help with FACS. We thank the Genome Technology Access Center at McDonnell Genome Institute at Washington University School of Medicine for help with snRNA-seq library construction and sequencing. We would also like to thank the Papouin lab at Washington University for use of NeuN, Iba1, and Sox9 antibodies. Lastly, we are grateful to members of the Creed and Samineni labs for their helpful discussion of the manuscript and experimental design.

## Author Contributions

V.K.S., M.C.C., and L.Y. conceptualized the project and formulated the overarching research goals. V.K.S., M.C.C., L.Y., and L.Z.F. developed and designed the experiments. V.C. performed NHP tissue dissections; V.K.S. and M.C.C. performed mouse tissue dissections. L.Y., H.H., Y.Z., and M.R.H. performed the sequencing experiments; L.Y. performed the sequencing analysis. L.Z.F, M.R.L., and C.S.X. performed FISH, immunohistochemistry, and analyzed the data; L.Y., H.H., L.Z.F., and M.R.L. generated the figures. L.Y., L.Z.F., V.C., V.K.S., and M.C.C. wrote the manuscript with input from all authors.

## Data and Materials Availability

All data needed to evaluate the conclusions in the paper are present in the paper and/or the Supplementary Materials. Raw and processed data of snRNA-seq experiments included in this study are deposited to the NCBI Gene Expression (GEO): GSE277465. Processed snRNA-seq data included in this study are available via Washington University Digital Commons Data@Becker with DOI: 10.17632/fyz3cn8r32.1.

---

**Fig. S1. Related to Fig. 1.**
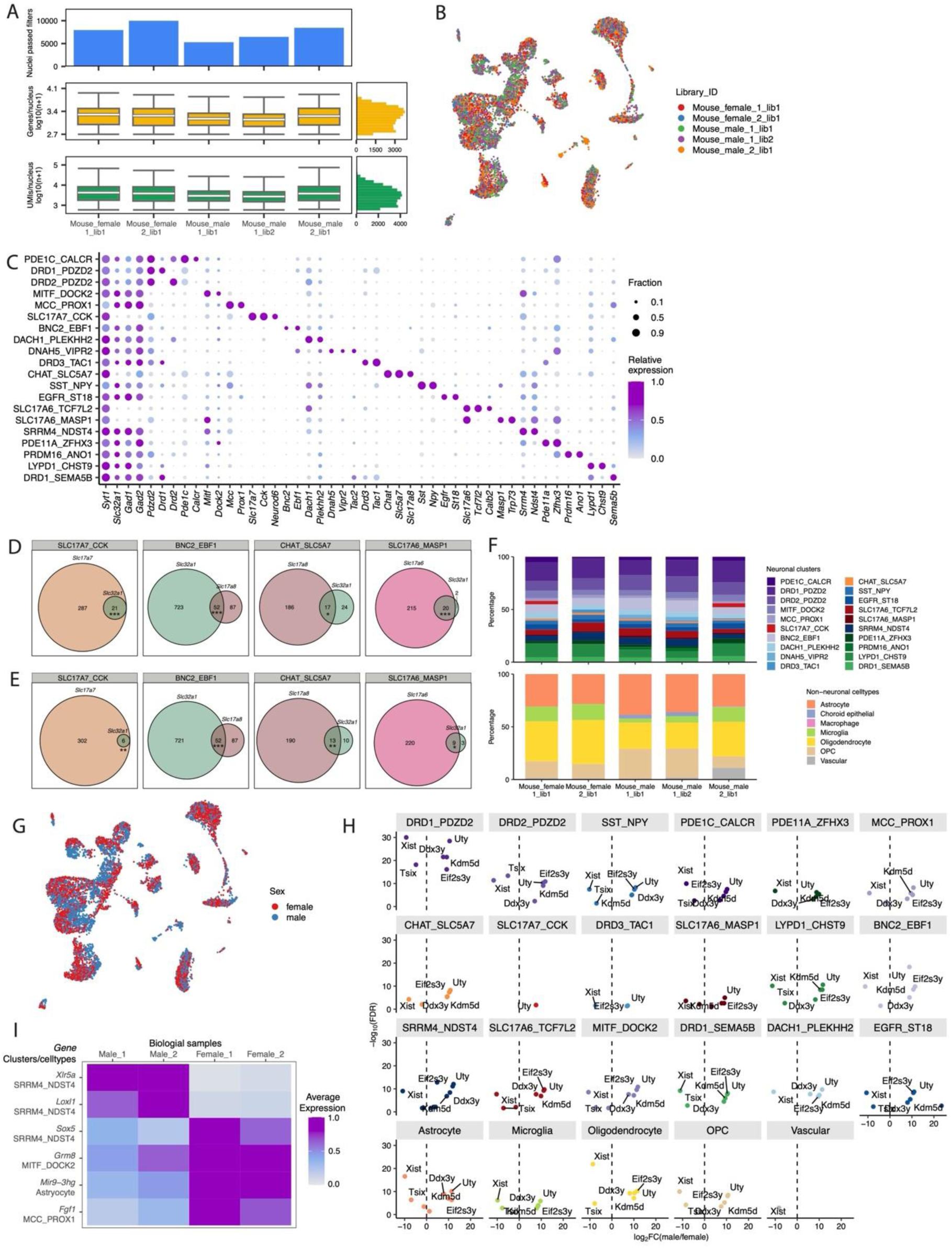
(**A**) Mouse snRNA-seq library metrics. The top row displays the number of nuclei passed quality control. The middle row displays box plots of the number of genes per nucleus (log10 transformed) and the bottom row displays the number of UMIs per nucleus. Boxes indicate quartiles and whiskers are 1.5-times the interquartile range (Q1-Q3). The median is a white line inside each box. The distribution is aggregated across all samples and displayed on the horizontal histogram. (**B**) UMAPs of 2,000 downsampled mouse nuclei per library. Nuclei are colored by libraries. (**C**) Dot plot displaying the expression of marker genes for mouse neuronal clusters. Dot size denotes the fraction of nuclei expressing a marker gene (>0 counts), and color denotes the relative expression of a gene in each cluster (calculated as the mean expression of a gene relative to the highest mean expression of that gene across all clusters). (**D**) Venn diagrams showing the nuclei in individual neuronal clusters co-expressing GABAergic and glutamatergic marker genes based on the original snRNA-seq counts. Numbers denote nuclei count in each group. Hypergeometric tests were performed to estimate the significance of overlap (SLC17A7_CCK: *p* = 5.2e-08, BNC2_EBF1: *p* = 2e-08, CHAT_SLC5A7: *p* = 0.02, CLS17A6_MASP1: *p* = 1.8e-06). (**E**) Venn diagram showing the nuclei in individual neuronal clusters co-expressing GABAergic and glutamatergic marker genes based on the snRNA-seq counts with ambient RNA contamination correction using soupX (*88*). Hypergeometric tests (SLC17A7_CCK: *p* =0.009, BNC2_EBF1: *p* = 1.9e-08, CHAT_SLC5A7: *p* = 0.002, CLS17A6_MASP1: *p* = 0.018). (**F**) Stacked bar plot showing the composition of neuronal clusters (top) and non-neuronal cell types (bottom) in each mouse snRNA-seq library. (**G**) UMAP of 5,000 downsampled mouse nuclei per sex. Nuclei are colored by sex. (**H**) Volcano plots of differentially expressed genes between nuclei from male (avg_log2FC > 0, FDR < 0.05) and female (avg_log2FC < 0, FDR < 0.05) in individual mouse cell types with at least 30 nuclei from each sex. (**I**) Heatmap showing the expression of six genes differentially expressed (FDR<0.05, pseudobulk DE analysis) between male and female mouse samples in respective neuronal clusters annotated on the left.

**Fig. S2. Related to Fig. 2.**
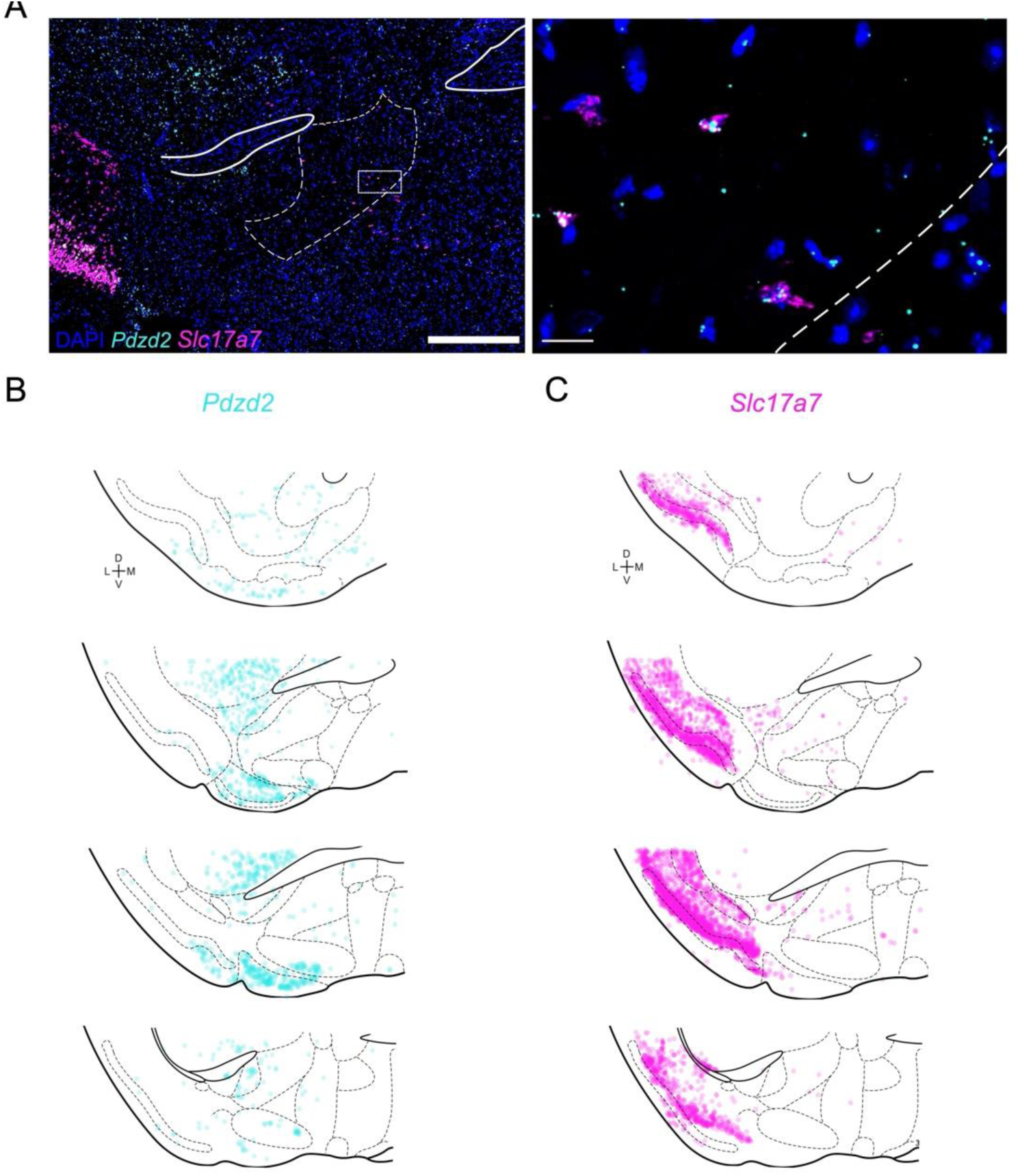
(**A**) Representative image of *Pdzd2 (cyan) and Slc17a7* (magenta) expression in the basal forebrain. Scale bars = 500 µm (left); 20 µm (right). (**B-C**) Fluorescent *in situ* hybridization data confirms the expression of *Pdzd2* (B) and *Slc17a7* (C) across the anteroposterior extent of the VP, and adjacent sub-commissural structures with biological replicates (n=3) overlaid.

**Fig. S3. Related to Fig. 2.**
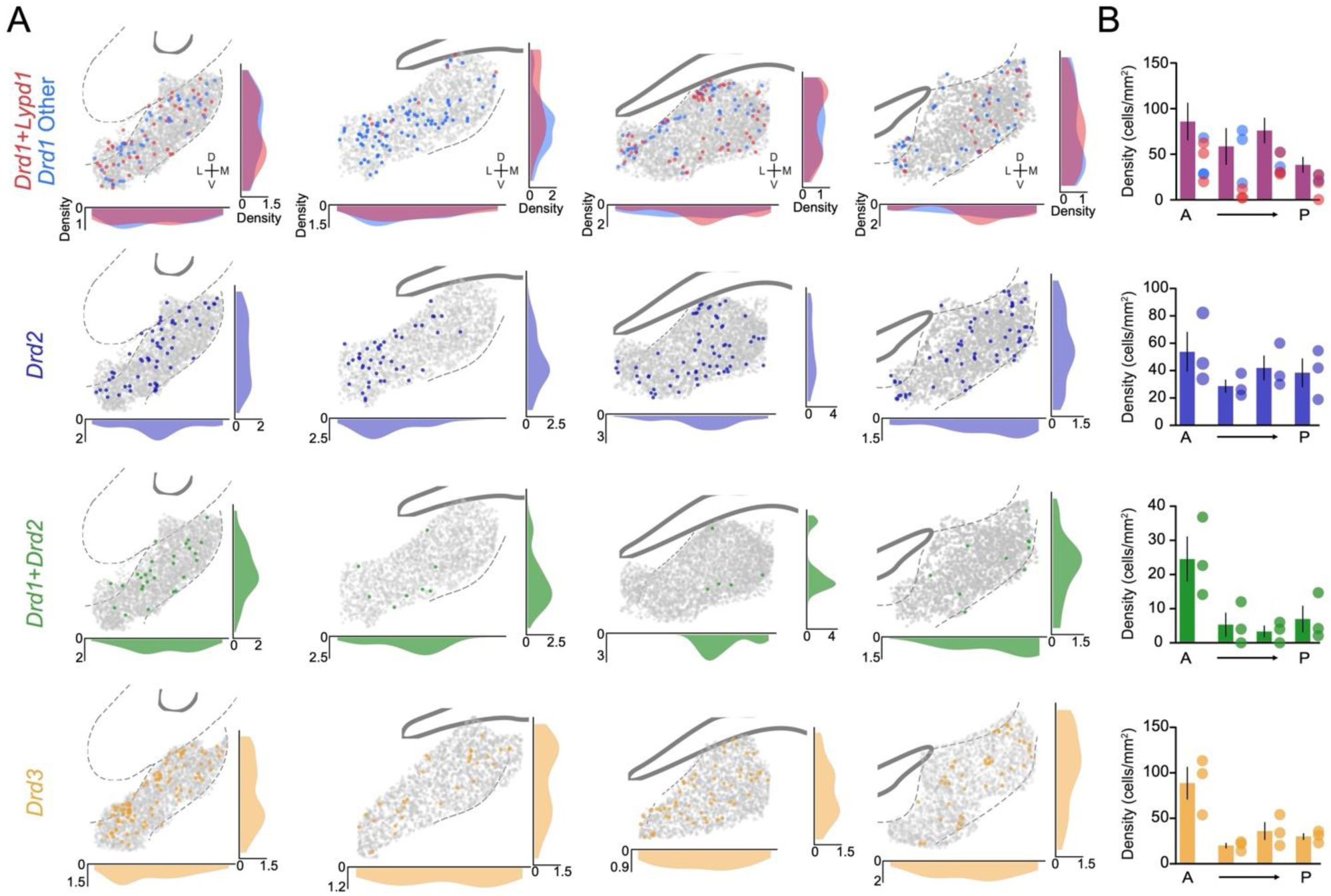
Spatial maps of dopaminoceptive cells over four levels of the VP as determined by HiPlex FISH. (**A**) Colorized dots correspond to cells that expressed probes that were hybridized to *Drd1*, *Drd2*, *Drd3*, and *Lypd1* mRNA. Density plots depict the relative distribution of cells over the mediolateral and dorsoventral extent of the VP. (**B**) Bar graphs indicating the density (cells/mm2) of individual clusters over the anteroposterior axis of the VP. Dots to the right of each bar represent one biological sample (n=3).

**Fig. S4.**
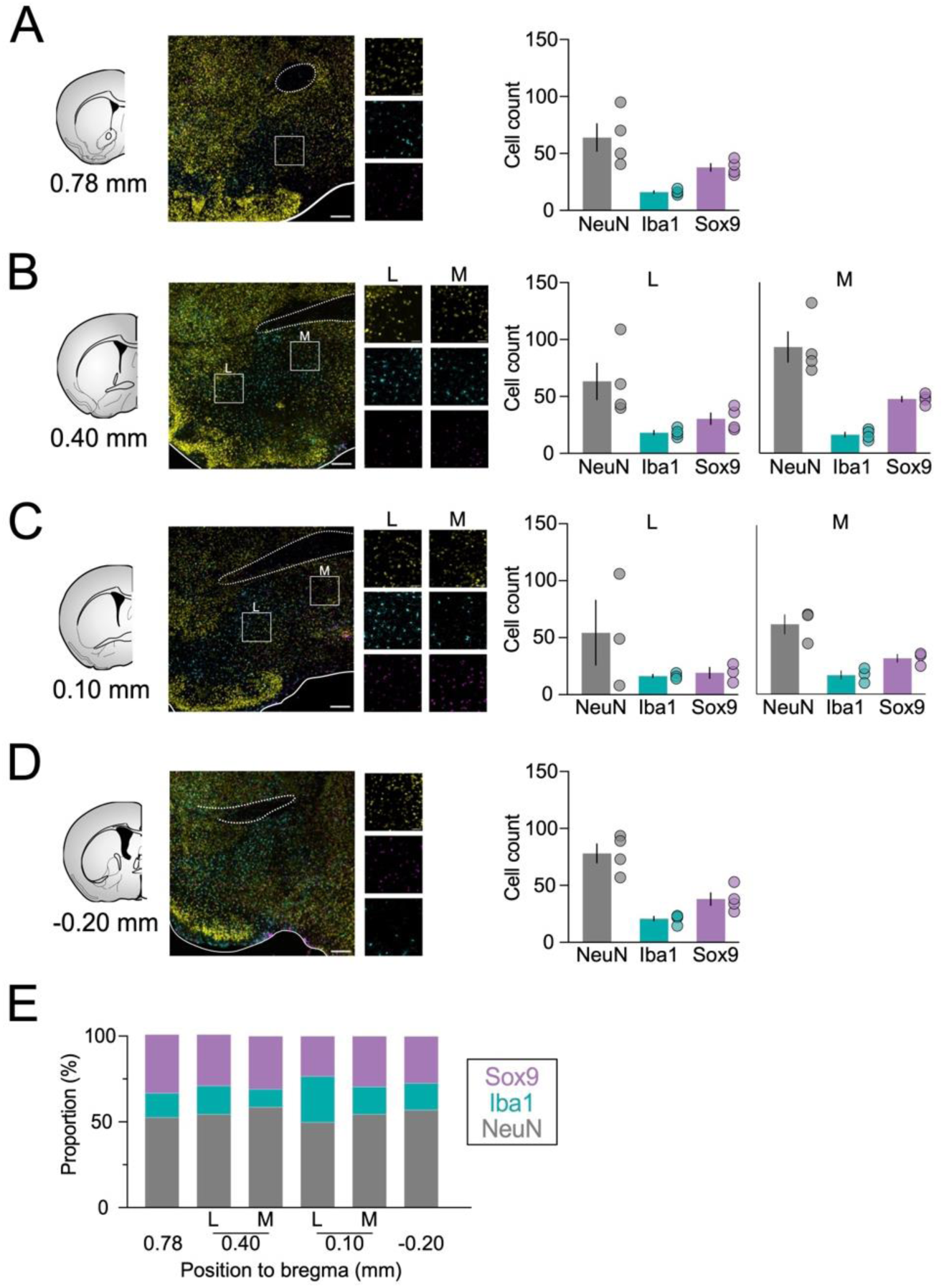
Distribution of neuronal and non-neuronal cell types in the VP of mice – Related to Fig. 2. (**A-D**) (Left) Representative images of NeuN (yellow), Iba1 (cyan), and Sox9 (magenta) immunohistochemical staining over four levels, spanning the anterior-posterior extent of the VP. Low magnification image scale bar = 200 µm; high magnification image scale bar = 50 µm. (Right) The number of neurons (NeuN-positive), microglia (Iba1-positive), and astrocytes (Sox9) spatially distributed across the VP. (**E**) Stacked bar plot showing the relative proportion of neurons, microglia, and astrocytes at each defined level of the VP.

**Fig. S5. Related to Fig. 4.**
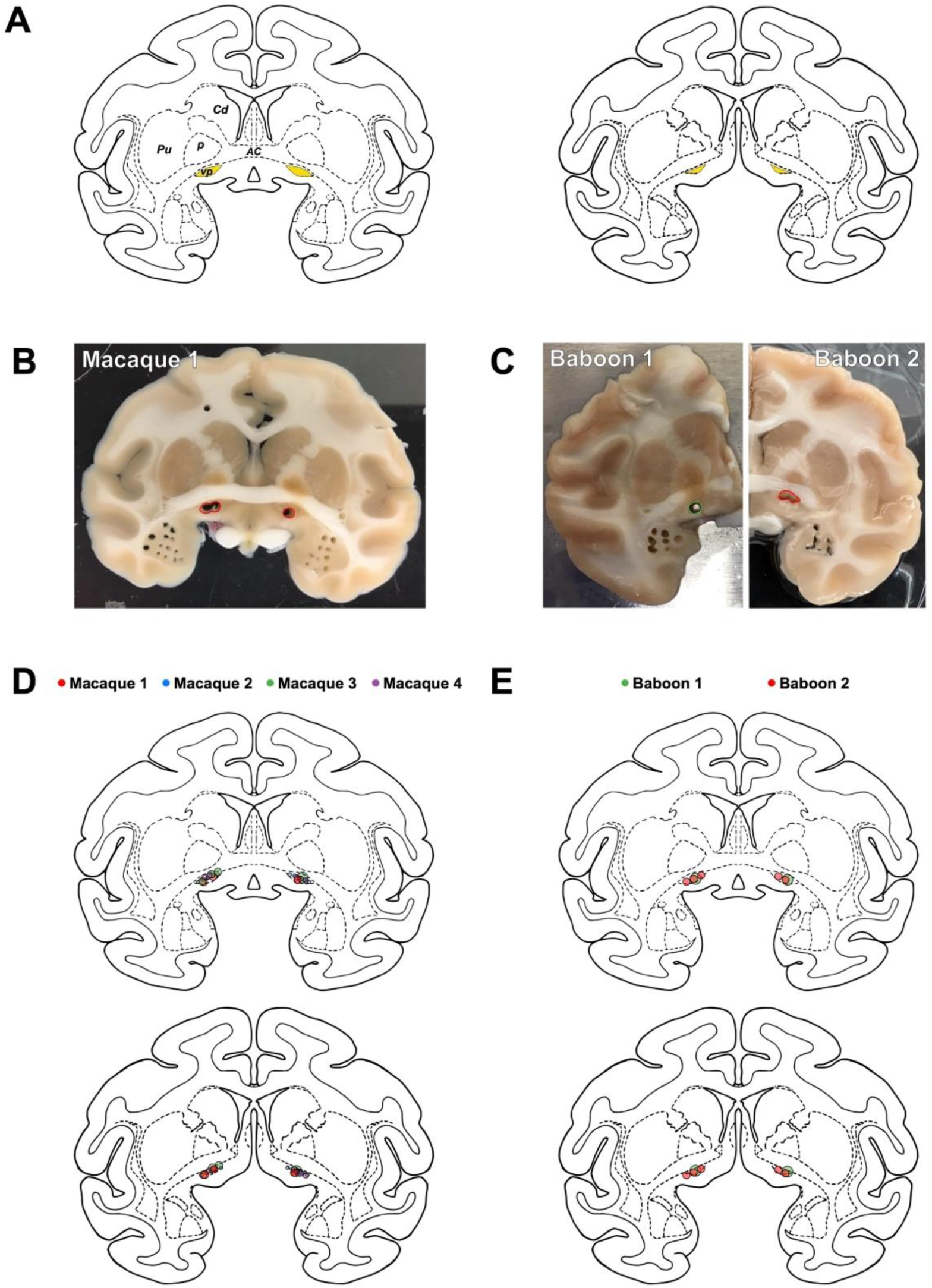
(**A**) Coronal sections of the NIMH Laboratory of Neuropsychology Rhesus Macaque “RED” Symmetrical Brain Atlas taken at levels +17 to +18 mm from ear bar zero depicting the location of the VP. (**B-C**) Representative images of coronal brain sections taken from macaque (B) and baboon (C); location of tissue punches is highlighted. (**D-E**) Composite schematic of VP tissue punches taken from each of 4 macaques (D) and 2 baboons (E).

**Fig. S6. Related to Fig. 4.**
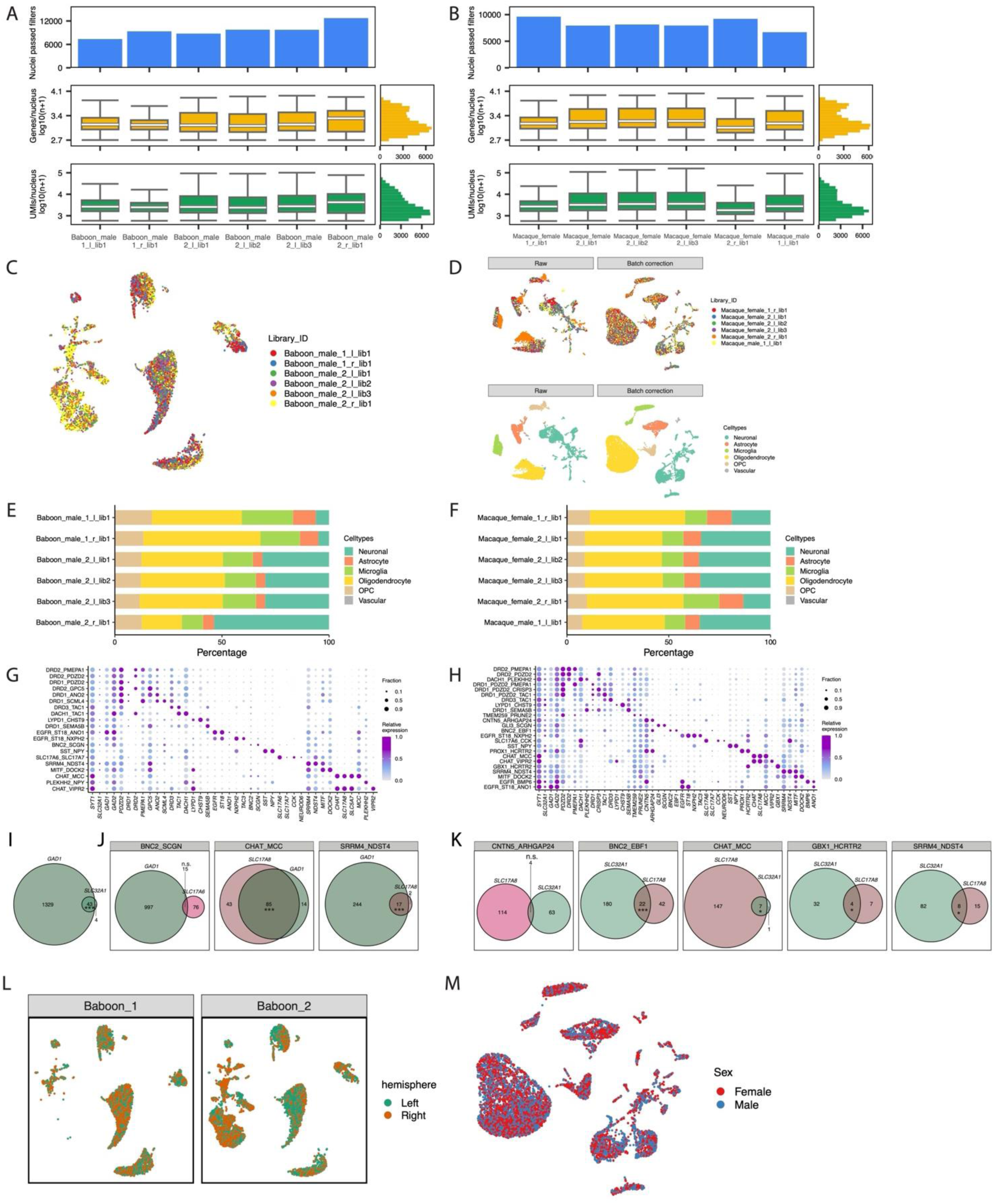
(**A-B**) snRNA-seq library metrics for baboon (A), and macaque (B). The top row displays the number of nuclei passed quality control. The middle row displays box plots of the number of genes per nucleus (log10 transformed) and the bottom row displays the number of UMIs per nucleus. Boxes indicate quartiles and whiskers are 1.5-times the interquartile range (Q1-Q3). The median is a white line inside each box. The distribution is aggregated across all samples and displayed on the horizontal histogram. (**C**) UMAPs of 2,000 downsampled baboon nuclei per library. Nuclei are colored by libraries. (**D**) Batch correction for the macaque dataset. 2,000 nuclei per library were randomly downsampled and shown on the UMAPs pre- (left) and post-batch correction (right). Nuclei are colored by libraries (top) or celltypes (bottom). (**E-F**) Stacked bar plot showing the composition of cell types in each baboon (E), and macaque (F) snRNA-seq library. (**G-H**) Dot plot displaying the expression of marker genes for baboon (G) and macaque (H) neuronal clusters. Dot size denotes the fraction of nuclei expressing a marker gene (>0 counts), and color denotes the relative expression of a gene in each cluster (calculated as the mean expression of a gene relative to the highest mean expression of that gene across all clusters). (**I**) Venn diagrams showing the baboon nuclei from neuronal clusters BNC2_SCGN, CHAT_MCC, and SRRM4_NDST4 co-expressing GABAergic marker genes *GAD1* and *SLC32A1*. Numbers denote nuclei count in each group. Hypergeometric test was performed to estimate the significance of overlap (*p* = 8e-18). (**J**) Venn diagrams showing nuclei from individual baboon neuronal clusters co-expressing GABAergic and glutamatergic marker genes. Hypergeometric tests (BNC2_SCGN: *p* = 0.99, CHAT_MCC: *p* = 2e-29, and SRRM4_NDST4: *p* = 6.3e-05). (**K**) Venn diagrams showing nuclei from individual macaque neuronal clusters co-expressing GABAergic and glutamatergic marker genes. Hypergeometric tests (CNTN5_ARHGAP24: *p* = 0.99, BNC2_EBF1: *p* = 8.7e-06, CHAT_MCC: *p* = 0.03, GBX1_HCRTR2: *p* = 0.05, and SRRM4_NDST4: *p* =0.02). (**L**) UMAPs of 5,000 downsampled baboon nuclei per hemisphere per animal. Nuclei are colored by the origin of the hemisphere. (**M**) UMAP of 5,000 downsampled macaque nuclei per sex. Nuclei are colored by sex.

**Fig. S7. Related to Fig. 5.**
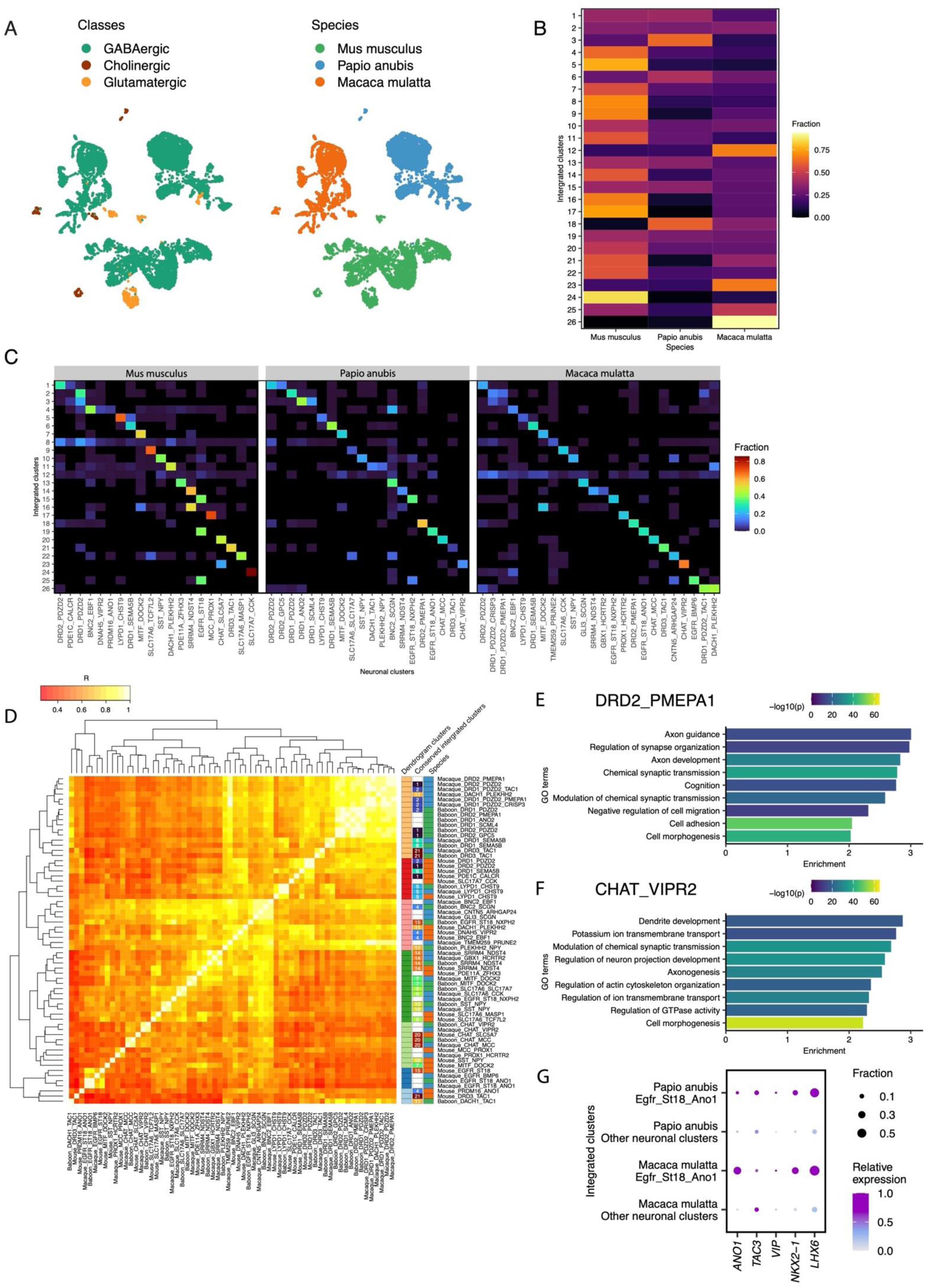
(**A**) UMAPs showing the clustering of VP neuronal nuclei across species without integration. 3,000 neuronal nuclei per species are downsampled from mouse, baboon, and macaque. Nuclei are colored by neuronal cell types (top left), species (top right), and integrated clusters from the harmonized atlas (bottom). (**B**) Heatmap showing the fraction of nuclei in each integrated cluster by species. (**C**) Heatmap showing the fraction of nuclei in each integrated cluster by each neuronal cluster identified by independent analysis of each species. (**D**) Heatmap showing the Pearson’s correlation coefficients of the expression of conserved marker genes (Log2FC>0.5, FDR<0.05 in all species) across individual neuronal clusters identified by independent analysis of each species. Neuronal clusters are ordered based on the hierarchical clustering of the expression of conserved marker genes used in the heatmap. (E and F) Significant Gene Ontology terms enriched in shared marker genes (Log2FC>0.5, FDR<0.05 in both species) of DRD1_PMEPA1 and CHAT_VIPR2 clusters distinct in NHPs. (G) Dot plot displaying the expression of select CGE-and MGE-specific genes in nuclei from EGFR_ST18_ANO1 cluster and other neuronal clusters in baboon and macaque.

**Fig. S8. Related to Fig. 5.**
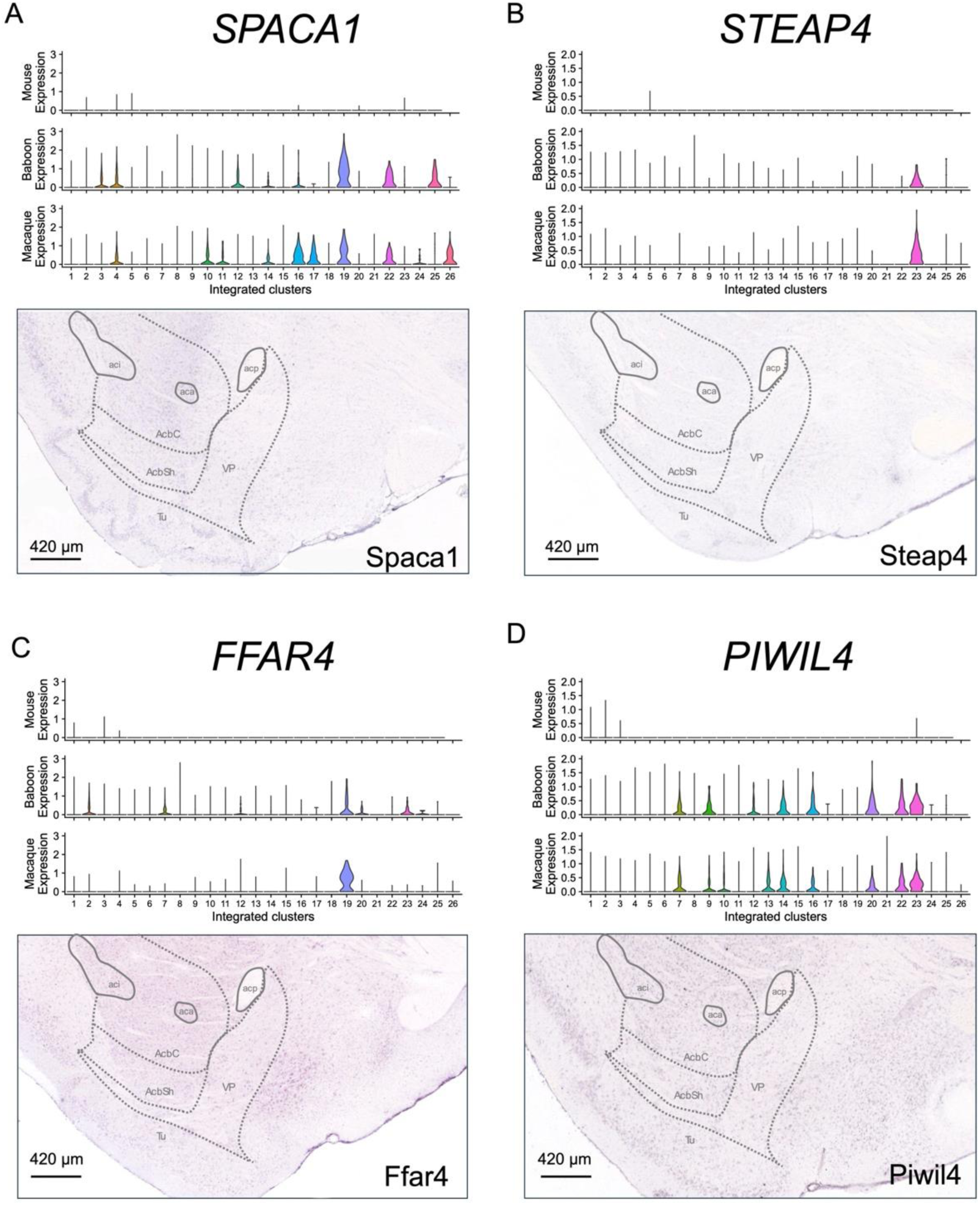
(**A-D**) Publicly available in situ hybridization data confirms the lack of expression of NHP-specific ventral pallidal genes in murine VP. Candidate genes SPACA1 (A), STEAP4 (B), FFAR4 (C), and PIWIL4 (D) were selected for representative enrichment in macaque and baboon VP subclusters. Top: Violin plots showing relative expression of the gene of interest in mouse (top), macaque (mid), and baboon (bottom). Bottom: In situ hybridization images from Allen Brain Atlas showing lack of expression of homologous genes in rodent VP. All sections show the sagittal plane at 1.2 mm from midline, brain regions are aligned to that atlas of Paxinos and Watson. Allen Mouse Brain Atlas, https://mouse.brain-map.org/experiment/show/74988286.

**Table S1. Metadata and sequencing metrics for individual snRNA-seq libraries**

File: Table S1_sample_metadata.xlsx

**Table S2. Marker genes for individual cell types and neuronal clusters in mouse snRNA-seq data**

File: Table S2_Mouse marker genes.xlsx

Differential expression analysis was done using FindAllMarkers() in Seurat comparing nuclei in one celltype/neuronal cluster to all other nuclei/neuronal nuclei. Genes with FDR < 0.05 were reported.

**Table S3. Differentially expressed genes in individual mouse cell types/neuronal clusters between sex**

File: Table S3_Mouse DE genes by sex.xlsx

pseudobulk counts for each neuronal cluster or non-neuronal cell type were generated by aggregating counts from nuclei of the same biological sample using AggregateExpression() in Seurat. Differential expression analysis was done using DESeq2 V1.44.0 in R by comparing counts of biological samples between sex (n = 2 for each sex). Genes with FDR < 0.05 were reported.

**Table S4. Marker genes for individual cell types and neuronal clusters in each NHP**

File: Table S4_NHP marker genes.xlsx

Differential expression analysis was done using FindAllMarkers() in Seurat comparing nuclei in one celltype/neuronal cluster to all other nuclei/neuronal nuclei in each species. Genes with FDR < 0.05 were reported.

**Table S5. Neuronal cluster correspondence across species in the harmonized atlas**

File: Table S5_cluster correspondence across species.xlsx

To facilitate accurate comparative analysis across species, snRNA-seq data from Baboon, macaque, and mouse were integrated and jointly clustered using Seurat (Methods). To establish correspondence between cross-species integrated clusters and mouse, macaque, and baboon clusters, the table shows each integrated cluster and its corresponding neuronal clusters from each species, which were generated independently by the clustering approach. A neuronal cluster was assigned to its integrated cluster if at least 35% of nuclei from that neuronal cluster were mapped to the corresponding integrated cluster.

**Table S6. Gene symbols of the genes mentioned in this study**

File: Table S6_GeneAbbrevs.xlsx

